# Mapping PTBP splicing in human brain identifies targets for therapeutic splice switching including *SYNGAP1*

**DOI:** 10.1101/2022.07.15.500219

**Authors:** Jennine M. Dawicki-McKenna, Alex J. Felix, Elisa A. Waxman, Congsheng Cheng, Defne A. Amado, Paul T. Ranum, Alexey Bogush, Lea V. Dungan, Elizabeth A. Heller, Deborah L. French, Beverly L. Davidson, Benjamin L. Prosser

**Affiliations:** Department of Physiology, Pennsylvania Muscle Institute, University of Pennsylvania Perelman School of Medicine; Center for Cellular and Molecular Therapeutics, Children’s Hospital of Philadelphia; Department of Systems Pharmacology and Translational Therapeutics, University of Pennsylvania Perelman School of Medicine; Department of Pathology and Laboratory Medicine, University of Pennsylvania Perelman School of Medicine

## Abstract

Alternative splicing of neuronal genes is controlled in part by the coordinated action of the polypyrimidine tract binding proteins (PTBP1 and PTBP2). While PTBP1 is ubiquitously expressed, PTBP2 is predominantly neuronal, controlling the expression of such targets as *DLG4*, which encodes PSD95, a protein important in synaptic function whose deficiency causes neurodevelopmental disorders. Here, we fully define the PTBP2 footprint in the human transcriptome using both human brain tissue and neurons derived from human induced pluripotent stem cells (iPSC-neurons). We identify direct PTBP2 binding sites and define PTBP2-dependent alternative splicing events, finding novel targets such as *STXBP1* and *SYNGAP1*, which are synaptic genes also associated with neurodevelopmental disorders. The resultant PTBP2 binding and splicing maps were used to test if PTBP2 binding could be manipulated to increase gene expression in PTBP-targeted genes that cause disease when haploinsufficient. We find that PTBP2 binding to *SYNGAP1* mRNA promotes alternative splicing and non-sense mediated decay. Antisense oligonucleotides that disrupt PTBP binding sites on *SYNGAP1* redirect splicing and increase gene and protein expression. Collectively, our data provide a comprehensive view of PTBP2-dependent alternative splicing in human neurons and human cerebral cortex, guiding the development of novel therapeutic tools that may benefit a range of neurodevelopmental disorders.

## Introduction

Alternative splicing (AS) of precursor mRNA (pre-mRNA) is an essential mechanism for post-transcriptional gene diversification and regulation. There is a particularly high frequency of AS in the brain, where it is required for all aspects of nervous system development and function. Concurrently, aberrant AS is implicated in multiple neurological disorders (for reviews, see^1–3^).

Therapeutic targeting of AS with antisense oligonucleotides (ASOs) that redirect splicing has demonstrated clinical potential for treating neurological disorders^4^. ASOs are short, single-stranded nucleic acid analogs that take advantage of Watson–Crick base pairing to target RNA molecules. Depending on their chemistry, ASO binding can result in target gene degradation or modulation of RNA processing (for review, see^5^). Since AS is controlled by trans-acting RNA-binding proteins (RBPs) that promote or repress splicing, steric-blocking ASOs that disrupt the interaction between these proteins and their target pre-mRNA can redirect AS to therapeutic benefit^6,7^. Spinraza® is a prominent example of a splice-switching ASO that exerts therapeutic benefit in the central nervous system disorder spinal muscular atrophy (SMA). This ASO binds to the *SMN2* pre-mRNA to disrupt a splice silencing RBP, in turn promoting *SMN2* exon 7 inclusion and augmented SMN protein expression, improving disease trajectory^8,9^.

AS of neuronal genes is controlled by the coordinated action of several prominent RBPs, which include the polypyrimidine tract binding proteins (PTBP1 and PTBP2). PTBP1 and PTBP2 are structurally similar and bind overlapping RNA targets yet differ by their cell type expression patterns. PTBP1 is broadly expressed across cell types but largely absent from neurons, while PTBP2 is predominantly neuronal (also referred to as “nPTB”). PTBP2 is required for neuron development and survival, and functions in part to suppress adult splicing patterns to control the temporal regulation of neuronal maturation^10–12^. Analysis of differentially expressed transcripts upon PTBP2 ablation suggests preferential regulation of targets involved in pre- and post-synaptic assembly and synaptic transmission^11^.

To identify binding sites of PTBPs, cross-linking immunoprecipitation followed by RNA sequencing (CLIP-seq) has been used in non-neuronal cells for PTBP1^13^ and mouse embryonic cortical tissue for PTBP2^12^. Yet to identify direct targets of PTBP2 splicing in the brain, transcriptome-wide PTBP2 binding must be combined with differential splicing analysis upon PTBP2 manipulation. Neither PTBP2 binding nor splicing has been assessed in human neurons nor post-embryonic brain tissue.

Here we determined the direct targets of PTBP2-dependent AS in the human brain. To achieve this, we performed CLIP-seq analysis of PTBP2 binding in both human cortical tissue and human neurons derived from induced pluripotent stem cells (iPSC-neurons), and we combined this with splicing analysis following PTBP2 depletion in iPSC-neurons **(Fig 1a)**. From this analysis we identified several genes that are alternatively spliced and repressed by PTBP2, and which cause human disease when reduced in expression. These negatively regulated genes represent potential targets for antisense splice switching for therapeutic upregulation. We performed extensive follow-up on one such gene, *SYNGAP1*, where variants lead to reduced expression (haploinsufficiency) and a phenotypically broad neurodevelopmental disorder^14–16^. We utilized PTBP2 binding and splicing maps to guide ASO disruption of PTBP2 binding to *SYNGAP1*, effectively redirecting splicing to increase gene expression. Together, this work provides the first comprehensive map of PTBP2-dependent AS in human neurons and cortical tissue and identifies targets for ASO manipulation that may offer therapeutic potential.

**Figure 1.**
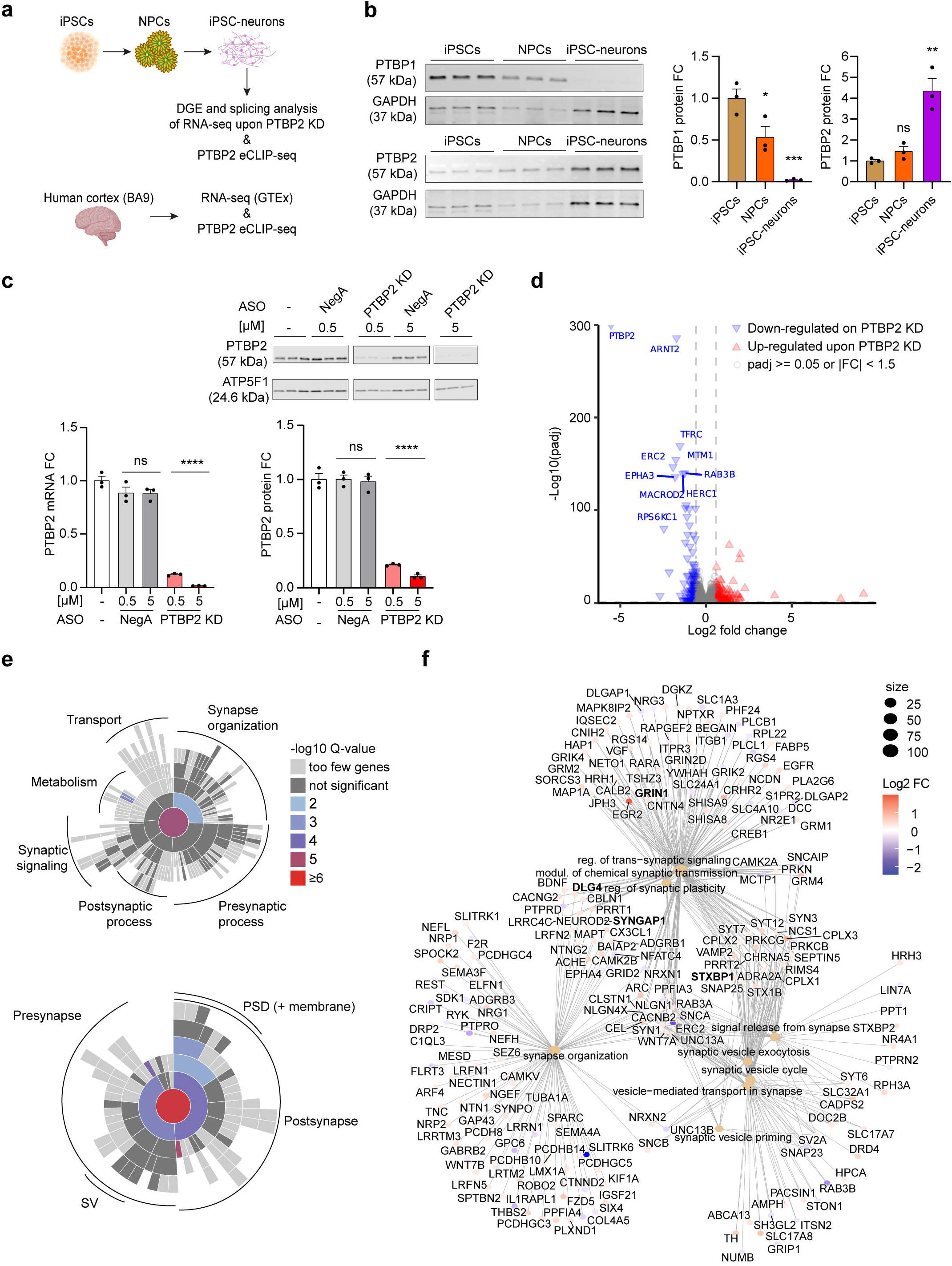
Differential gene expression upon PTBP2 depletion in iPSC-neurons. **(a)** Cartoon schematic of experimental design. **(b)** Western blot of PTBP protein levels at different stages of neuronal maturation. **(c)** Validation of PTBP2 depletion using “gapmer” ASOs in iPSC-neurons. (Left) qPCR and (Top and bottom right) Western blot of PTBP2. NegA, negative control gapmer. **(d)** Volcano plot of differential gene expression comparing untreated and PTBP2 KD iPSC-neurons. **(e)** SynGO enrichment analysis of curated synaptic genes differentially expressed upon PTBP2 KD (padj < 0.05, |fold-change| >= 1.15) relative to a background set of brain-expressed genes represented as a sunburst plot. (Top) Biological Process, 224 genes. (Bottom) Cellular Component, 271 genes. **(f)** Category netplot (clusterProfiler) of the top synapse-associated GO terms overrepresented upon PTBP2 KD (Biological Process, PTBP2 KD vs. untreated iPSC-neurons) relative to a background of all genes evaluated. Color indicates fold-changes of differentially expressed synapse-associated genes (padj < 0.05, |fold-change| >= 1.15). Size indicates number of genes in term. *n* = 3 biological replicates for iPSC-neurons RNA-seq data sets. In **b** and **c**, data are represented as mean values ± SEM. All data points represent independent biological replicates. b and c (*n* = 3). **b**, one-way ANOVA with Dunnett’s multiple comparison test vs iPSCs. **c**, one-way ANOVA with Dunnett’s multiple comparison test vs mock-treated cells (−). ns p > 0.05, *p < 0.05, **p < 0.01 and ***p< 0.001.

## Results

### PTBP2 differentially regulates synaptic genes

To identify targets of PTBP2-dependent AS, we aimed to deplete PTBP2 in cortical excitatory neurons with functional synaptic activity that were derived from human iPSCs (see Materials and Methods; **Supplementary Fig. 1a-c; Supplementary Movie 1**). We first characterized PTBP1 and PTBP2 expression over the course of iPSC differentiation into neurons. Consistent with previous literature on the developmental regulation of PTBP isoforms^17,18^, we noted high levels of PTBP1 in undifferentiated iPSCs that decreased in neuronal progenitor cells (NPCs) and was minimally detectable in iPSC-neurons. Conversely, PTBP2 expression increased during differentiation and exhibited the highest expression in iPSC-neurons (**Fig. 1b**). To deplete PTBP2, we utilized locked-nucleic acid (LNA) modified ASOs (“gapmers”) to trigger RNase H1-dependent degradation of PTBP2 mRNA. After initial screening in HEK293T cells (**Supplementary Fig. 1d**), the most effective gapmer (PTBP2_7, hereafter referred to as PTBP2 KD) was examined for dose-dependent knockdown of PTBP2 in iPSC-neurons 7 days after gymnotic delivery. PTBP2 KD produced a robust reduction of PTBP2 mRNA and protein expression, with PTBP2 protein levels reduced by >90% (**Fig. 1c)**.

We next performed RNA sequencing upon PTBP2 KD in iPSC-neurons. Principal component analysis indicated that PTBP2 KD drove the majority of variance in this data set, with tight clustering between biological replicates and between the two negative control groups **(Supplementary Fig. 1e)**. Differential gene expression (DGE) analysis identified significantly up- and down-regulated genes and confirmed PTBP2 as the most significantly decreased gene upon PTBP2 depletion (**Fig. 1d, full dataset in Supplementary Data 1)**. Gene ontology analysis of biological processes indicated prominent alterations in genes involved in cell cycle regulation, DNA repair, synaptic transmission and plasticity (**Supplementary Fig. 1f)**. Synaptic ontology terms curated by SynGO^19^ demonstrated enrichment in genes differentially regulated by PTBP2 relative to a background set of brain-expressed genes (**Fig. 1e**). Within the synapse groups, 304 genes associated with synaptic transmission or plasticity were differentially regulated upon PTBP2 KD, including well-known targets of PTBP2 such as *DLG4* that encodes the major synaptic scaffolding protein (PSD-95), as well as novel targets such as *GRIN1, STXBP1* and *SYNGAP1*, which encode prominent regulators of pre-and post-synaptic function (**Fig. 1f)**.

### PTBP2 alters splicing and expression of various disease-causing genes

We next evaluated transcriptome-wide AS upon PTBP2 KD using replicate multivariate analysis of transcript splicing (rMATS)^20^ (**Fig. 2a)**. Using a false discovery rate (FDR) <0.05 and inclusion level difference of 5%, 1389 genes were differentially spliced upon PTBP2 depletion. The use of rMATS identified 5 distinct AS events: exon skipping, alternative 5’ and 3’ splice sites (SS), mutually exclusive splicing events, and retained introns. For each event type, >100 genes were differentially spliced upon PTBP2-depletion, with exon skipping the most predominantly regulated event **(Fig. 2a, GSE206660 rMATS Supplementary file)**. Both inclusion and exclusion of AS events were observed upon PTBP2 KD, although across all 5 types of events the depletion of PTBP2 was more likely to promote event exclusion. These findings indicate that the endogenous presence of PTBP2 preferentially promotes the inclusion of AS events in human iPSC-neurons.

**Figure 2.**
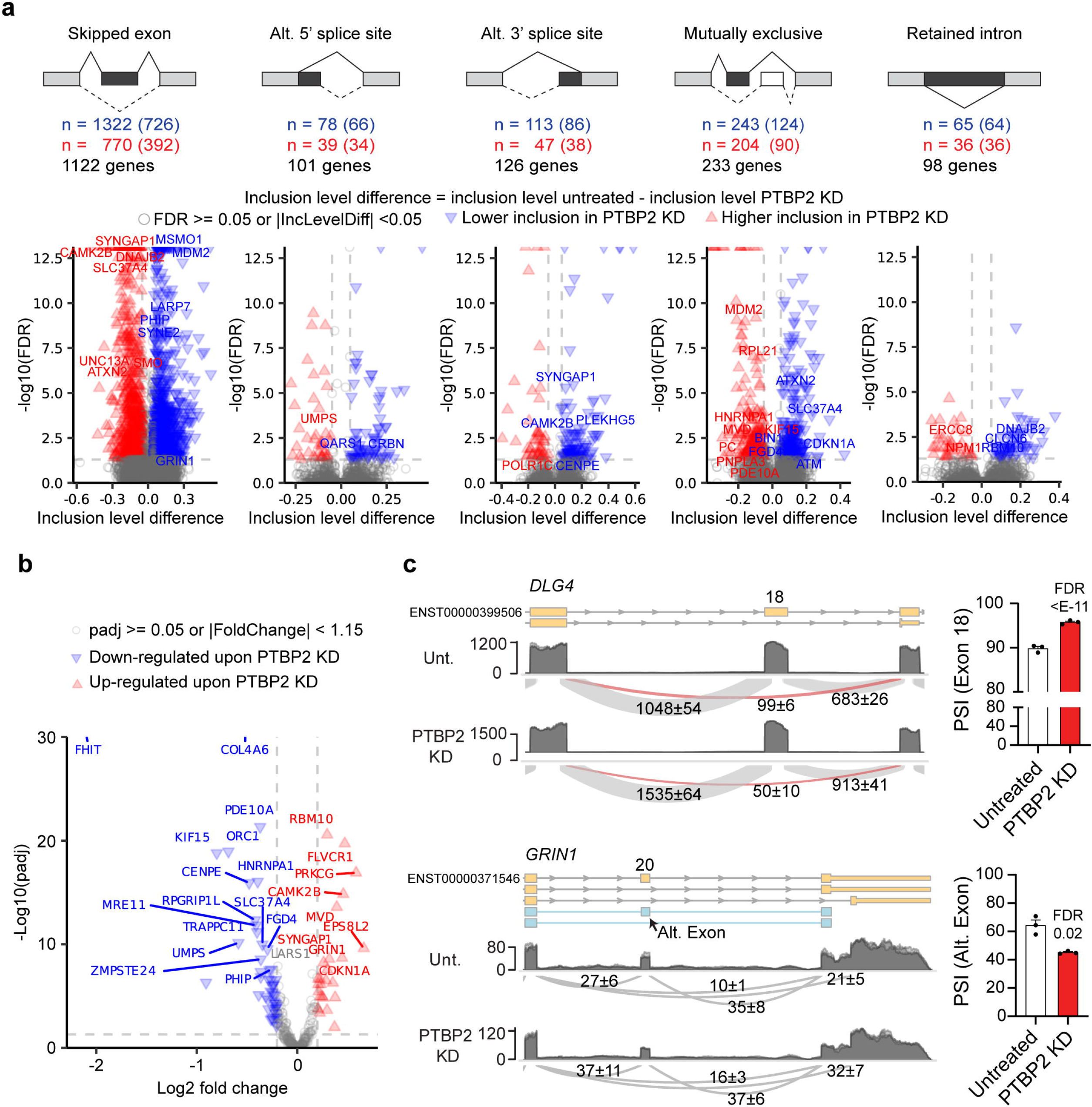
Differential splicing analysis following PTBP2 depletion in iPSC-neurons. **(a)** (Top) Types of alternative splicing (AS) events, number of alternative events detected (number of nonoverlapping events in parentheses), and number of genes having at least one AS event for each type in PTBP2 KD vs. untreated iPSC-neurons. Fractional inclusion level difference is inclusion level untreated - inclusion level PTBP2 KD and refers to inclusion of the darkly shaded element. (Bottom) Volcano plot for each alternative splicing type. Orphanet genes with the smallest FDRs are annotated. **(b)** Volcano plot of differential gene expression for the subset of Orphanet genes that are differentially spliced in PTBP2 KD vs. untreated iPSC-neurons (FDR < 0.05, |inclusion level difference| >= 0.05). (**c)** Representative AS events for *DLG4* (top, encoding PSD-95) and *GRIN1* (bottom) shown as sashimi plots (left; numbers indicate the number of reads spanning junctions±SD) with replicates overlaid; at right is the percent spliced in (PSI) for the AS event detected by rMATS. Statistical significance determined in rMATS by a likelihood-ratio test at a cutoff of 1% difference. *n* = 3 biological replicates for iPSC-neurons RNA-seq data sets.

Given our interest in neurodevelopmental disorders, we cross-referenced this list of alternatively spliced transcripts regulated by PTBP2 with the Orphanet database of disease-causing genes. Upon PTBP2 depletion, 342 genetic etiologies were differentially spliced including *GRIN1, MVD, DNM1, CAMK2B, HNRNPA1, CTNND1* and others (**Fig. 2a, Supplementary Data 2**). Of these Orphanet genes, *SYNGAP1*, the genetic cause of SYNGAP1-associated intellectual disability, showed the most significant alternative 3’ splice site regulation as well as robust exon skipping upon PTBP2 KD (**Fig. 2a**). Intriguingly, AS of *SYNGAP1* was recently demonstrated to promote nonsense-mediated decay (NMD) and restrict *SYNGAP1* expression in HEK293 cells^6^, but neither the mechanism of this AS nor its neuronal impacts were explored.

Of the differentially spliced and disease-causing genes, 72 (including *SYNGAP1* and *GRIN1*) were also differentially expressed at the mRNA level upon PTBP2 KD, suggesting that PTBP2-dependent AS regulates the splicing and expression of several disease-causing genes (**Fig. 2b, c, Supplementary Data 2)**. Figure 2C shows a closer examination of the AS of *DLG4*, an event well documented in other systems that we use as a positive control^17^, and *GRIN1*, which has not been explored. Consistent with previous literature, we found that PTBP2 promotes the exclusion of *DLG4* exon 18 (i.e. KD increases inclusion), which triggers NMD and restricts expression of PSD-95^17^. For *GRIN1*, rMATS identified significant PTBP2-dependent AS to a previously unannotated alternative exon that introduces a frameshift from the canonical transcript and is associated with reduced expression (**Fig. 2b, c**).

### Mapping PTBP2 binding in the human brain

To increase confidence in PTBP2-dependent AS targets and identify direct sites of PTBP2 binding, we next performed PTBP2 CLIP-seq in iPSC-neurons and human cortical brain tissue (Brodmann Area 9). Transcriptome-wide CLIP-seq analysis and CLAM-mediated peak calling^21^ identified 49,875 and 57,162 high-confidence PTBP2 binding sites (bin size of 50 nucleotides) distributed across 8822 and 7869 genes in the human cortex and iPSC-neurons, respectively **(GSE206650 Supplementary files)**. Consistent with the above splicing analysis, CLIP-seq identified PTBP2 binding sites just upstream of the alternatively-spliced exon 18 of *DLG4* in both iPSC-neurons and human cortex (**Fig. 3a**). PTBP2 binding sites were predominantly intronic (72%), yet when normalized to the number of nucleotides, PTBP2 binding was more evenly distributed across 5’UTRs, coding regions, and 3’UTRs **(Fig. 3b)**. Motif analysis in both the human brain and iPSC-neurons indicated CUCUCU as the most enriched sequence observed in PTBP2 binding sites, consistent with previous identification of the preferred binding sequence for PTBP proteins^22^ **(Fig. 3c)**. Gene ontology analysis of genes with PTBP2 peak calls demonstrated a predominant overrepresentation of synaptic terms (**Fig. 3d, Supplementary Fig. 2a)**, and synaptic ontology terms curated by SynGO^19^ were enriched in genes with PTBP2 peak calls relative to a background set of brain-expressed genes (**Fig. 3e; Supplementary Fig. 2b**).

**Figure 3.**
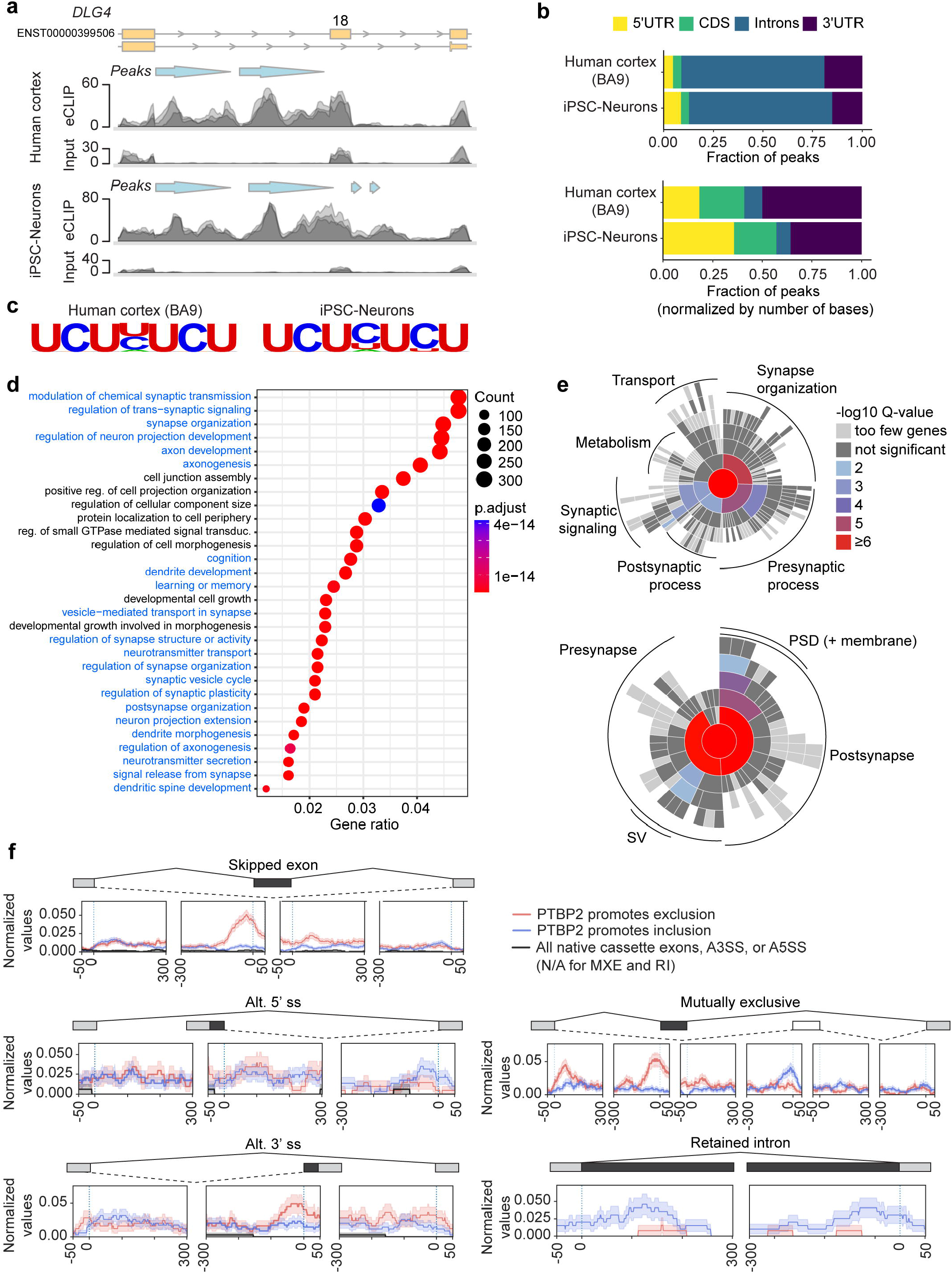
PTBP2 CLIP-seq in iPSC-neurons and human cortex. (**a**) Representative PTBP2 eCLIP read density relative to size-matched input (replicates overlaid) on *DLG4* (encoding PSD-95) showing called PTBP2 binding peaks (light blue arrows) at similar locations in human cortex (top) and iPSC-neurons (bottom) in the intronic region upstream of Exon 18, a known PTBP2 splicing target. **(b)** Fraction of PTBP2 CLIP-seq peaks by genomic region for iPSC-neurons and human cortex, prior to or following normalization for number of bases in the respective region type. **(c)** Motif enrichment analysis for PTBP2 CLIP-seq. Top ranked motifs are shown for human cortex (p-val 1e-2761, 28.6% of targets) and iPSC-neurons (p-val 1e-677, 19.3% of targets). **(d)** Dotplot (clusterProfiler) showing the top 30 categories from GO enrichment analysis (Biological Process) of PTBP2 CLIP-seq peaks in human cortex (BA9). Gene ratio is number of genes with peak calls relative to total genes in GO group (count). Synapse-related terms are highlighted in blue. **(e)** SynGO enrichment analysis of PTBP2 CLIP-seq peaks in human cortex (BA9) relative to a background set of brain-expressed genes represented as a sunburst plot. (Top) Biological Process, 623 genes. (Bottom) Cellular Component, 755 genes. **(f)** RBP-Maps-mediated positional analysis of PTBP2 peak calls relative to AS events identified as differentially spliced by rMATS upon PTBP2 KD in iPSC-neurons. Red indicates a higher inclusion level of the darkly shaded element upon PTBP2-KD, hence it suggests that endogenous PTBP2 promotes exclusion of the darkly-shaded element (n = 1217, 203, 245, 553, and 127 for SE, A5SS, A3SS, MXE, and RI events, respectively). Blue indicates the opposite (n = 1806, 286, 342, 657, and 196 for SE, A5SS, A3SS, MXE, and RI events, respectively). *n* = 3 biological replicates for PTBP2 CLIP-seq and size-matched input controls.

By cross-referencing our differentially spliced transcripts upon PTBP2 KD with positional data from CLIP-seq analysis in iPSC-neurons, we identified 316 AS events across 237 genes that demonstrated PTBP2 binding within 350 nucleotides of the AS event. We classify these as high-confidence, direct AS events mediated by proximal PTBP2-binding (**Supplementary Data 2)**.

We also performed a more detailed positional analysis using RBP-Maps^23^ to interrogate the transcriptome-wide relationship between sites of PTBP2 binding and the differential splicing detected upon PTBP2 KD **(Fig. 3f)**. This analysis indicates that PTBP2 binding within ∼100nt upstream of a cassette exon promotes skipping (exclusion) of that exon, as is the case for *DLG4* exon 18 (**Fig. 2c and Fig. 3a**). A similar relationship was observed for PTBP2 binding upstream and promoting the exclusion of the proximal exon in mutually exclusive AS events. While more variability is evident in the analysis of less frequently detected events (alternative 5’ and 3’ splice sites, retained introns), PTBP2 binding within ∼100nt in the upstream intron was also associated with the exclusion of 3’ alternative splice sites, while PTBP2 binding proximal to a downstream exon promoted inclusion of alternative 5’ splicing upstream. Intriguingly, PTBP2 binding was also found to promote intron retention, regardless of whether this binding occurred at the 5’ or 3’ end of the retained intron **(Fig. 3f)**.

To examine the conservation of PTBP2 binding and splicing between iPSC-neurons and the human cortex, we evaluated the overlap in PTBP2 binding at three resolutions. At the gene level, the overlap in PTBP2 binding was 37% (**Supplementary Fig. 2c**). At the 50-nt resolution used for CLAM-mediated peak calling, 12.8% of PTBP2 binding sites were shared between iPSC-neurons and adult human cortex (**Supplementary Fig. 2c**). Finally, we evaluated PTBP2 binding overlap within 350 nucleotides proximal to each detected AS event in iPSC-neurons (**Supplementary Fig. 2d**). For example, if PTPB2 promoted exclusion of a cassette exon (the most predominant event in our dataset) and binding was observed in iPSC-neurons, it was observed in the same window in the human cortex for 38% of detected events. These data indicate the persistence of PTBP2-dependent AS for fine-tuning neuronal gene programs into adulthood.

### PTBP2 promotes the incorporation of a premature termination codon in SYNGAP1

Of the 342 Orphanet genes differentially spliced upon PTBP2 KD, 75 also showed PTBP2-binding proximal to one or more AS events (**Supplementary Data 2)**. This permits the evaluation of whether PTBP2-dependent splicing could be targeted for therapeutic benefit for these genetic etiologies. We investigated AS of *SYNGAP1*, a gene where variants cause haploinsufficiency resulting in phenotypically diverse neurodevelopmental disorders characterized by seizures, global delay, and autism^14–16^. By comparing RNA sequencing data from adult human cortex with iPSC-neurons, we identified conserved AS events on *SYNGAP1* (**Supplementary Fig. 3a)**. This includes the well-studied 5’ alternative transcriptional start sites (TSS; leads to “A” and “B” N-terminal SYNGAP1 variants) and AS of the final 3’ exon, which creates SYNGAP1 C-terminal variants α1 and α2^24^. Also exhibited in both datasets is the 3’AS event at Exon 11, a partial inclusion of exon 14, and inclusion of a predicted “poison” exon after Exon 18 (alternate exon 19, A.E. 19x) **(Supplementary Fig. 3a)**. rMATS splicing analysis indicates that PTBP2 promotes the inclusion of the 3’ AS of Exon 11 **(Fig. 4a, b, left)**, as well as the exclusion of Exon 14 **(Fig. 4a, b, middle)**, while not affecting the inclusion of A.E. 19x **(Fig. 4a, b, right)**.

**Figure 4.**
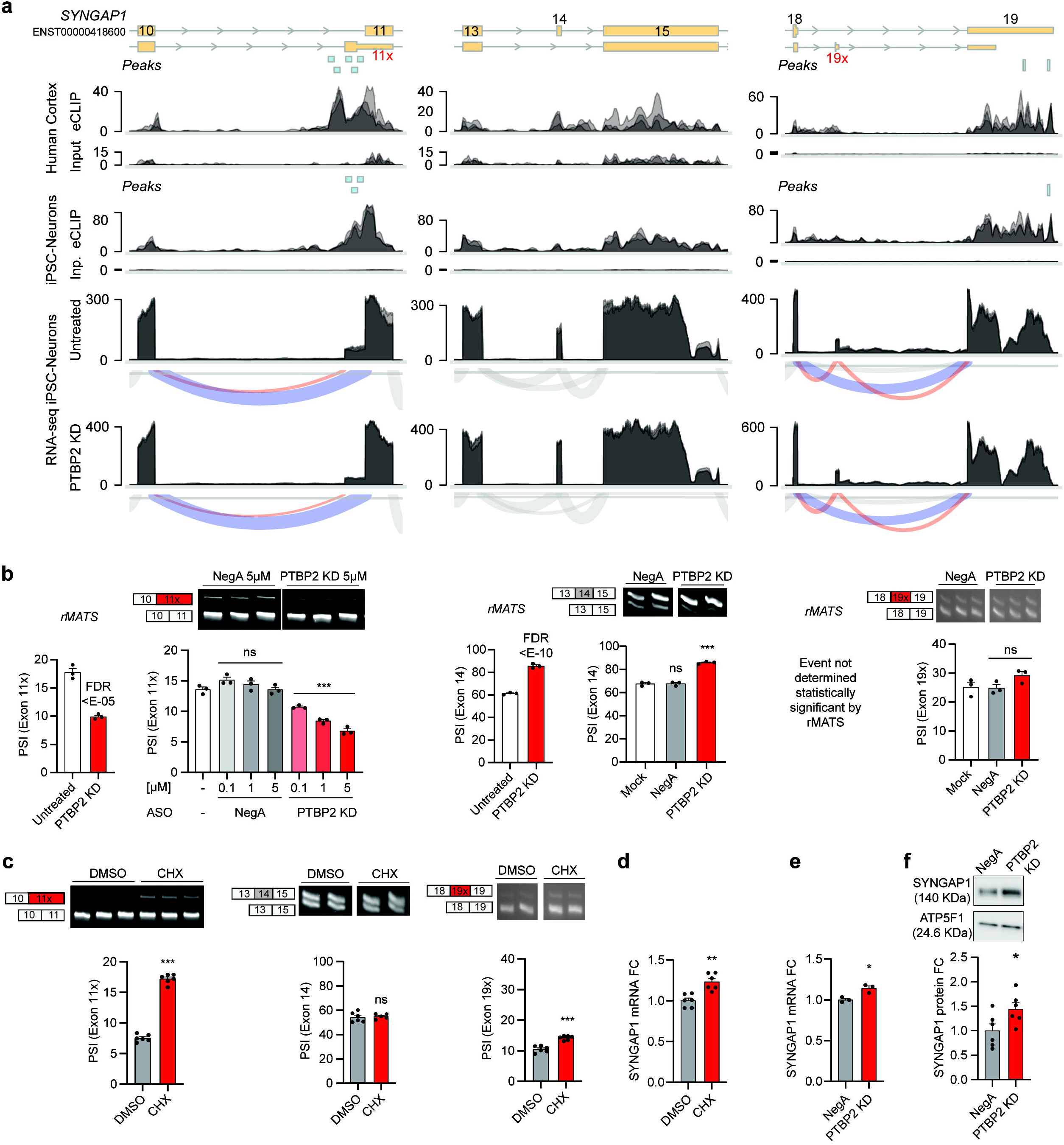
PTBP2 binds and promotes differential splicing and nonsense-mediated decay of *SYNGAP1*. (**a**) Zoom in for three regions of interest on *SYNGAP1* showing (top) gene model of alternative splicing event (ENST00000418600 is the dominant isoform in brain), followed by (middle) human cortex CLIP-seq (peaks, PTBP2 eCLIP read coverage, size-matched input read coverage), then iPSC-neurons CLIP-seq (as for cortex). Bottom two rows depict RNA-seq read coverage and sashimi plots for untreated control and PTBP2 KD. *n* = 3 biological replicates (overlaid) for human cortex CLIP-seq and iPSC-neurons RNA-seq and CLIP-seq data sets. **(b)** Quantification of changes in AS at each of the 3 above regions of interest (aligned in column) upon PTBP2 KD by rMATS (left, 5 µM, statistical significance determined in rMATS by a likelihood-ratio test at a cutoff of 1% difference) and RT-PCR splicing assays (right). Light shaded rectangles denote constitutive exons and red rectangles denote NMD-inducing AS events. **(c)** RT-PCR splicing assays of each region of interest upon CHX treatment to inhibit NMD. **(d)** Quantification (qPCR) of *SYNGAP1* mRNA fold change upon CHX treatment as measured across exon 16-17. **(e)** Quantification of *SYNGAP1* mRNA fold change upon PTBP2 KD. **(f)** SYNGAP1 protein expression by western blot upon PTBP2 KD. In **b-f**, data are represented as mean values ± SEM. All data points represent independent biological replicates. **b, e** (*n* = 3). **c** (*n* = 6, except *n* = 5 for CHX in middle panel). **d, f** (*n* = 6). In **b**, one-way ANOVA with Dunnett’s multiple comparison test vs mock-treated cells (−). In **c-f**, Student’s t-test. ns p > 0.05, *p < 0.05, **p < 0.01 and ***p< 0.001.

To confirm these findings, we designed RT-PCR splicing assays to assess PTBP2-dependent AS at each of these locations in human neurons. Consistent with rMATS analysis, PTBP2 KD led to a decrease in 3’AS of Exon 11, an increase in the inclusion of Exon 14, and had no significant effect on A.E. 19x **(Fig. 4b)**.

The 3’AS of Exon 11 and A.E. 19x are each predicted to introduce a premature termination codon that induces NMD, which has been demonstrated in non-neuronal cells for Exon 11^6^, while the inclusion of Exon 14 does not alter the reading frame. We evaluated each potential site as a target for NMD in iPSC-neurons by using the NMD inhibitor cycloheximide (CHX) and confirmed that NMD inhibition significantly increased 3’AS of Exon 11 **(Fig. 4c, left)**. CHX had a modest effect on A.E. 19 (**Fig. 4c, right**) while not affecting Exon 14 inclusion (**Fig. 4c, middle**). Consistent with relief from NMD, CHX treatment also increased total *SYNGAP1* mRNA expression (**Fig. 4d**). Further, PTBP2 KD and concomitant exclusion of the NMD-linked AS event resulted in increased SYNGAP1 mRNA and a 44 +/-14% increase in protein expression in human neurons **(Fig. 4e, f)**.

Taken together, the data show that PTBP2 binds directly to *SYNGAP1* (Fig. 4a, Supplementary Fig. 3b) to promote the inclusion of an alternative 3’ start site in Exon 11 causing NMD and repressed expression of SYNGAP1 in iPSC-neurons. As this AS event is also present in adult human cortex **(Supplementary Fig. 3a)**, disrupting PTBP2 binding to this region could be therapeutically beneficial in the context of SYNGAP1 haploinsufficiency. While targeting PTBP2 directly has limited therapeutic potential due to its numerous gene targets and pleiotropic actions (even on *SYNGAP1*), site-specific disruption using steric-blocking ASOs offers a precise approach.

### ASO disruption of PTBP binding improves SYNGAP1 splicing and expression

For high throughput screening of steric-blocking ASOs we utilized HEK293T cells, a cell type previously demonstrated to exhibit high levels of the 3’AS of *SYNGAP1* Exon 11^6^. We first examined whether PTBP also regulates this AS event in HEK293T cells as it does in neurons. Unlike in neurons, HEK293T cells predominantly express PTBP1, but PTBP paralogs share a consensus binding sequence and often have overlapping targets^25^. We thus utilized siRNA to knock down both PTBP1 and PTBP2 in HEK293T cells and examined *SYNGAP1* splicing. We first confirmed robust depletion of each isoform at the protein level **(Fig. 5a)** and found that PTBP1 depletion led to increased expression of PTBP2, as expected given the known downregulation of PTBP2 by PTBP1^26,27^.

**Figure 5.**
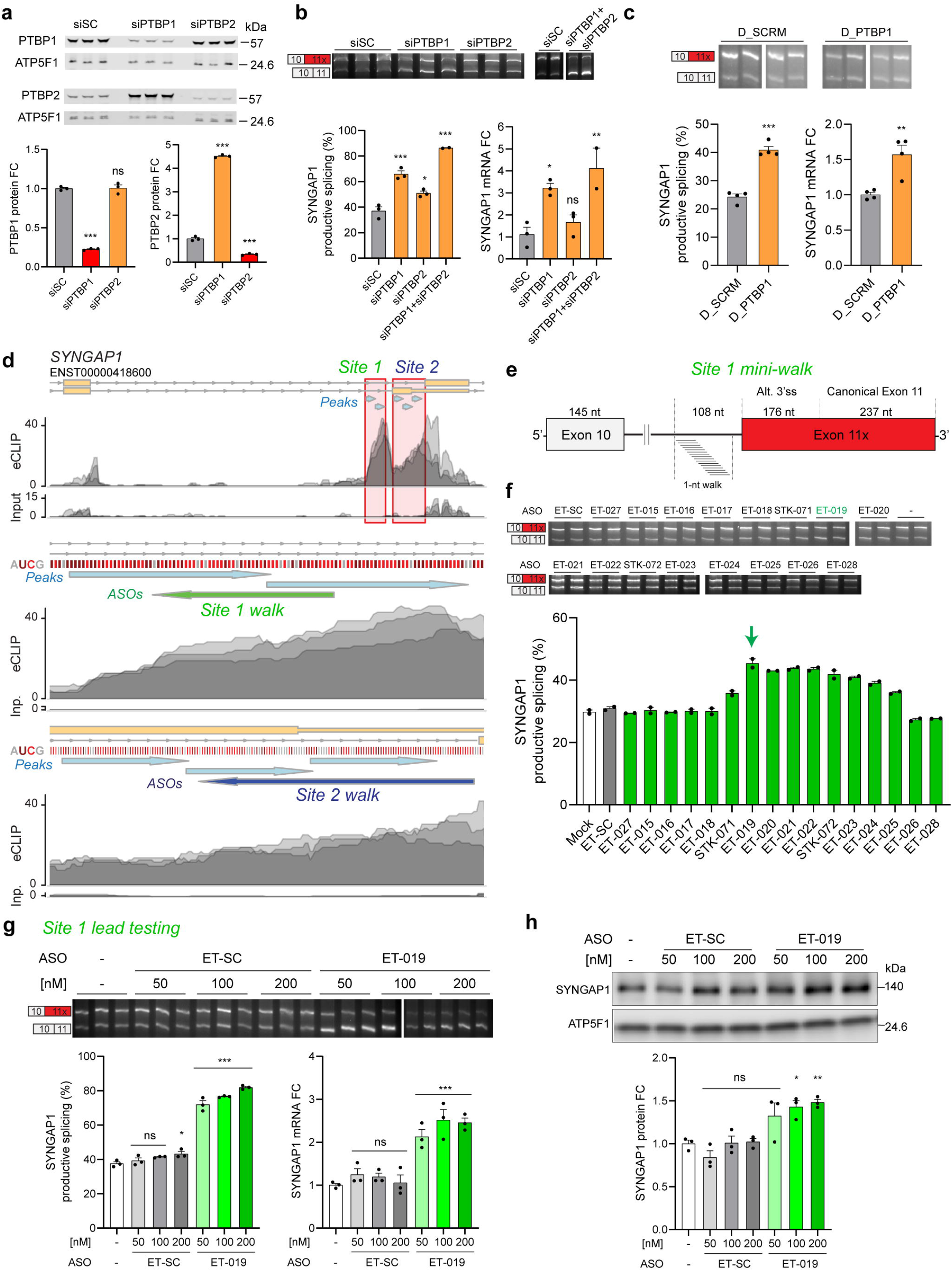
Disrupting PTBP binding in *SYNGAP1* intron 10 site upregulates SYNGAP1. (**a**) Western blot from HEK293T cells transfected for 48 h with siRNA against PTBP1 or PTBP2. siSC is a non-targeting control siRNA. (**b**) Top and left panel: RT-PCR from HEK293T cells transfected for 48 h with siRNA against PTBP1 or PTBP2 alone or in combination. Right panel: qPCR showing SYNGAP1 mRNA levels. (**c**) Top and left panel: RT-PCR from HEK293T cells transfected with 5 µM of PTBP decoy oligo for 48 h, including a non-targeting decoy (D_SCRM) as negative control. Right panel: qPCR showing SYNGAP1 mRNA levels. (**d**) Visualization of PTBP2 eCLIP-seq data from human cortex highlighting two highly enriched PTBP binding regions near SYNGAP1 exon 11: site 1 (green, intronic site) and site 2 (blue, NMD-inducing Exon 11x). Zoom-ins for both sites are provided including information about the nucleotide content (CU rich regions in red) and the location of the ASO walks. Light blue arrows indicate called peaks. (**e**) Scheme depicting 1-nt resolution ASO walk on SYNGAP1 site 1. The target region spans 33 nt of intronic sequence. Black lines denote introns, white rectangles denote constitutive exons and red rectangle denotes non-productive alternative exon (exon 11x). (**f**) RT-PCR from HEK293T cells transfected with 200 nM of ASO for 24 h, including a non-targeting ASO control (ET-SC) and no ASO control (Mock, -). (**g**) Top and left panel: RT-PCR from HEK293T cells transfected with increasing concentrations of lead ASO ET-019 and negative controls for 48 h. Right panel: qPCR showing SYNGAP1 mRNA levels. (**h**) Western blot from HEK293T cells transfected as in (**g**). Data are represented as mean values ± SEM. All data points represent independent biological replicates. **a, g** and **h** (*n* = 3). **b** (*n* = 3 except *n* = 2 for siPTBP1 + siPTBP2). **c** (*n* = 4). **f** (*n* = 2). In **a** and **b**, one-way ANOVA with Dunnett’s multiple comparison test vs siSC. In **c**, Student’s t-test. In **g** and **h**, one-way ANOVA with Dunnett’s multiple comparison test vs mock-treated cells (−). ns p > 0.05, *p < 0.05, **p < 0.01 and ***p< 0.001.

HEK293T cells exhibited a high percentage of the NMD-linked AS of *SYNGAP1* Exon 11 **(Fig. 5b)**, consistent with this event being increasingly excluded during neuronal maturation. Depletion of PTBP1 and to a lesser extent PTBP2 each led to the exclusion of this AS event and a proportional increase in the percentage of non-NMD, “productive” splicing of *SYNGAP1*, which was concomitant with increased *SYNGAP1* mRNA expression **(Fig. 5b)**. Dual depletion of PTBP1 and PTBP2 led to the greatest exclusion of the AS event and associated increase in *SYNGAP1* mRNA (**Fig. 5b**).

We next used an orthogonal approach to confirm PTBP-dependent regulation using an established PTBP decoy assay^28^. We transfected HEK293T cells with a high concentration (5 µM) of a decoy oligonucleotide containing 4 repeats of the PTBP consensus binding motif CUCUCU, which competes for PTBP occupancy of endogenous binding sites. Decoy transfection was sufficient to increase productive splicing of *SYNGAP1* at Exon 11 and *SYNGAP1* mRNA expression **(Fig. 5c)**. Further, there was high inclusion of this AS event in immortalized neuroblastoma cells (mouse N2a, human SHSY5Y) (**Supplementary Fig. 4a, b**), and gapmer-mediated PTBP1 depletion (**Supplementary Fig. 4a**) or ASO disruption of PTBP binding (**Supplementary Fig. 4b**) improved *SYNGAP1* productive splicing. Together with the above, these findings provide strong support that 1) PTBP binding promotes the inclusion of an alternative 3’ splice site in *SYNGAP1* Ex11 causing NMD and restricted expression across numerous cell types; 2) that both PTBP1 and PTBP2 can regulate this event; 3) the inclusion of this AS event decreases with neuronal maturity, concomitant with reduced levels of PTBP and increased *SYNGAP1* expression **(Supplementary Fig. 4c)**.

We next tested if disrupting PTBP binding improved productive *SYNGAP1* splicing using steric blocking ASOs. PTBP CLIP-seq analysis from both iPSC-neurons and the human brain pinpointed two PTBP binding regions near *SYNGAP1* Exon 11 to evaluate for therapeutic targeting (**Fig. 5d**). The first is a hot spot for PTBP enrichment and consensus binding sequences in Intron 10, approximately 100 nucleotides upstream from the alternative 3’ splice site, hereafter referred to as “Site 1”. The second is a highly enriched PTBP binding region that lies directly in the alternatively spliced-in region (“Site 2”).

Informatively, a recent 5 nucleotide spaced ASO “walk” found that several ASOs directed toward Site 1 elicited improvement in *SYNGAP1* productive splicing, although the mechanism for this was not interrogated^6^. We overlayed these results with our PTBP-binding maps and found that the effective ASOs indeed overlapped with PTBP binding sites. We refined this interrogation by performing a 1-nucleotide resolution ASO “mini-walk” in HEK293T cells (**Fig. 5e**) around the most promising region (near the previously identified STK-071) and found that ASOs spanning position -102 to -85 (ET-019) and -101 to -84 (ET-020) showed the most improvement in *SYNGAP1* productive splicing **(Fig. 5f)**. We also confirmed the effectiveness of ET-019 in SH-SY5Y cells (**Supplementary Fig. 4b**). In additional screenings to attempt optimization, modified versions of ET-019 containing extra nucleotides either in the 5’ or the 3’ end of the ASO did not improve productive splicing compared to ET-019 (**Supplementary Fig 4d**), and longer ASOs (22-mer) targeting the adjacent upstream region of ET-019 were also not effective (**Supplementary Fig. 4e**). In follow-up lead testing, we found ET-019 caused a dose-dependent improvement in SYNGAP1 productive splicing, mRNA, and protein expression **(Fig. 5g, h)**. These experiments identified ET-019 and ET-020 as the most promising ASOs for SYNGAP1 upregulation targeting site 1.

Our PTBP CLIP-seq data also directed us to interrogate site 2, which lies within the exonic region introduced by the alternative 3’ splice site and which has not been previously explored. A 5-nt resolution ASO walk around this region identified multiple candidates that could improve productive splicing of Exon 11 **(Fig. 6a)** and increase SYNGAP1 mRNA levels (**Supplementary Fig. 5a**). A follow-up on ET-061 showed increased expression of the productive transcript concomitant with robust improvements in *SYNGAP1* mRNA expression in a dose-dependent manner **(Supplementary Fig. 5b)**. As ET-061 was the most 5’ASO in the initial walk, we extended and refined this region upstream using an upstream mini-walk (**Fig. 6b**). Our data show that ASOs just upstream of ET-61 (ET-085 and ET-086) robustly improve both productive splicing (**Fig. 6b**) and total SYNGAP1 mRNA levels (**Fig. 6c**). Follow-up studies confirmed that ET-086 led to a dose-dependent increase in *SYNGAP1* productive splicing, mRNA, and protein expression (**Supplementary Fig. 5c, d)**.

**Figure 6.**
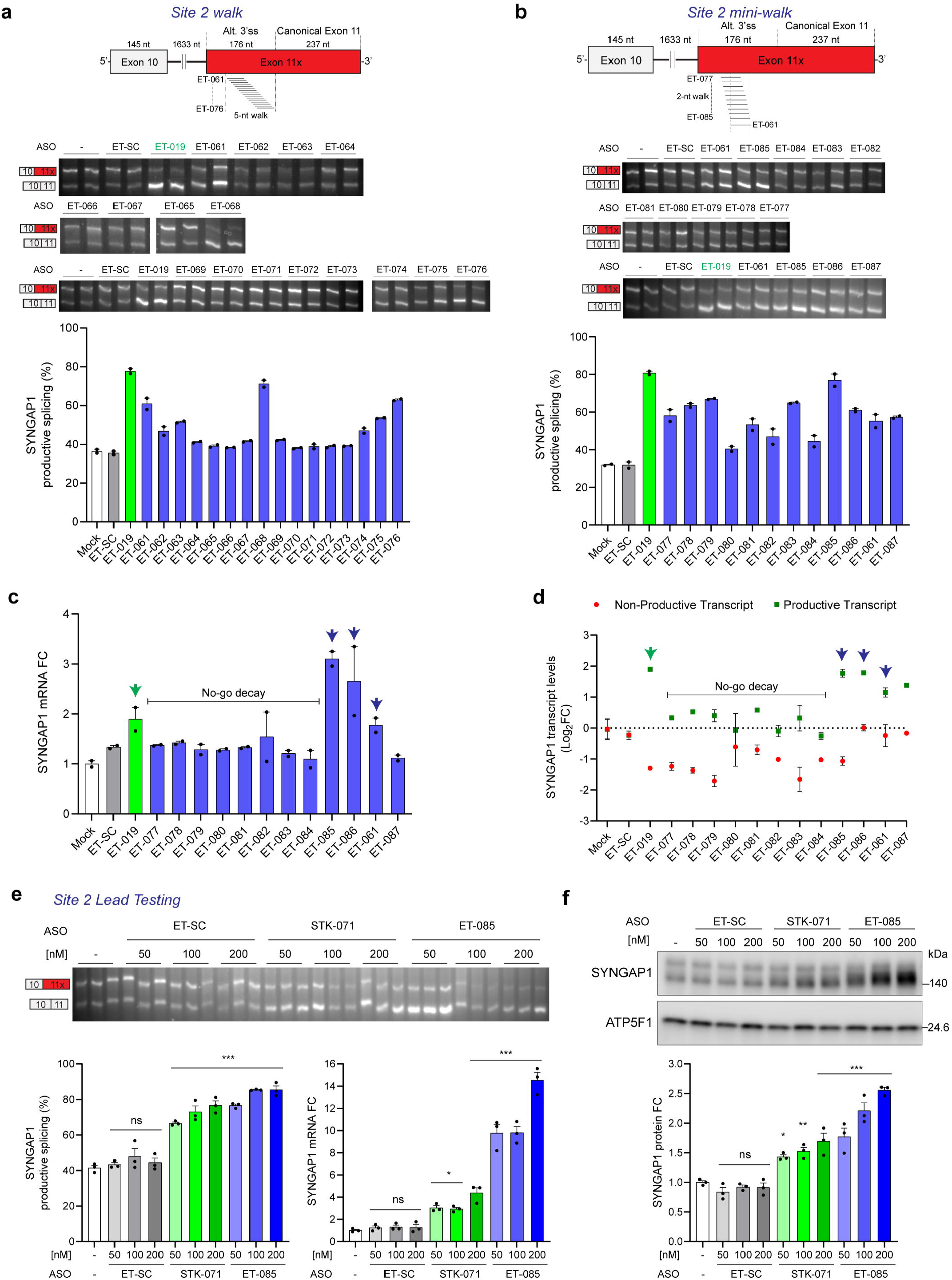
Disrupting PTBP binding in *SYNGAP1* exon 11x upregulates SYNGAP1. **(a)** Top, Scheme depicting the initial 5-nt resolution ASO walk. The target region spans 93 nt of non-productive 3’ss in *SYNGAP1*. Bottom, RT-PCR from HEK293T cells transfected with 100 nM of ASO for 24 h, including a positive control ASO targeting SYNGAP1 site 1 (ET-019), a non-targeting ASO control (ET-SC) and no ASO control (Mock, -). **(b)** Top, Scheme depicting a combined 2-nt and 1-nt resolution ASO walk. The target region spans 38 nt of non-productive 3’ss in SYNGAP1. Bottom, RT-PCR from HEK293T cells transfected with 100 nM of ASO for 24 h, including the same controls as in (a). **(c)** qPCR quantification of SYNGAP1 transcript levels from samples in (b). **(d)** Non-productive and productive transcript levels calculated from densitometric analysis of RT-PCR products from (b) and represented as log_2_FC values relative to Mock. Arrows indicate ASOs that increase *SYNGAP1* productive transcript and mRNA levels. Black line indicates ASOs that lead to no-go decay. **(e)** Top and left panel: RT-PCR from HEK293T cells transfected with increasing concentrations of STK-071, ET-085 and ET-SC for 48 h. Right panel: qPCR showing SYNGAP1 mRNA levels. **(f)** Western blot from HEK293T cells transfected as in (e). Data are represented as mean values ± SEM. All data points represent independent biological replicates. **a**-**d** (*n* = 2). **e, f** (*n* = 3). In **e** and **f**, one-way ANOVA with Dunnett’s multiple comparison test vs mock-treated cells (−). ns p > 0.05, *p < 0.05, **p < 0.01 and ***p< 0.001.

Intriguingly, several identified candidates (e.g. ET-079) improved productive splicing (**Fig. 6b**) but not *SYNGAP1* mRNA expression (**Fig. 6c**); ASOs that bound between +64 and +95 in the alternatively spliced-in region reduced expression of the non-productive transcript without a concomitant increase in the productively spliced transcript, as would be expected if a splice-switching oligo promoted utilization of an alternative splice site (**Fig. 6d**). These results are consistent with these ASOs binding to the non-productive transcript and triggering “no-go decay”, a mechanism whereby steric blocking oligos bound to coding sequences can disrupt ribosomal translation and induce transcript degradation in an RNase H1-independent manner^29^.

We noted that across (but not within) experiments the magnitude of the effect of a given ASO might vary, which was anecdotally linked to differences in cell plating density. As this prevented quantitative comparisons across different experiments, we directly compared our top-performing Site 1 and 2 oligos concurrently under identical conditions. We first evaluated the previously identified ASO targeting site 1 (STK-071) with the newly identified ASO targeting site 2 (ET-085) and found that each offers a dose-dependent improvement in SYNGAP1 productive splicing, mRNA expression, and protein expression (**Fig. 6e, f**), but with ET-085 showing dramatic upregulation at the mRNA level (to >10-fold) coupled with a greater increase in protein expression (2.5-fold). This site 2 targeting ASO offers comparable levels of SYNGAP1 upregulation at 4-fold lower concentration, which may increase its therapeutic potential by limiting oligo dosage and off-target effects.

A similarly designed experiment comparing multiple site 1 and 2 targeting ASOs revealed ET-019 and ET-085 as the most robust ASOs in terms of improving productive splicing and SYNGAP1 mRNA levels, respectively (**Supplementary Fig. 5e**). Of note, while improvements in productive splicing correlated with increased mRNA within both Site 1 and 2 targeting oligos, ASOs targeting Site 2 uniformly produced a larger increase in *SYNGAP1* mRNA relative to the change in productive splicing. This suggests additional - and incompletely understood - gain by targeting this region.

### ASO-mediated upregulation of SYNGAP1 in vivo

We next performed a pilot proof-of-concept study to examine whether Site 1 and 2 targeting ASOs can upregulate *SYNGAP1* in the postnatal brain *in vivo*. We designed homologous mouse-targeting ASOs equivalent to our top performing site 1 (ET-019) and site 2 (ET-085) oligos and performed intracerbroventricular injections (ICV) into the mouse brain at post-natal day 2 then harvested the brain 5 days later (**Supplementary Fig. 6a)**. We included STK-135, a splice-switching oligo demonstrated to potently bind and manipulate *SCN1A*^6^, as a positive control to demonstrate target engagement of a neuronal gene in our hands, and as a negative control ASO de-targeted from *SYNGAP1*. Consistently, we found that STK-135 potently improved productive splicing and expression levels of *SCN1A* (**Supplementary Fig 6b, c**), with no effect on *SYNGAP1* productive splicing or mRNA expression **(Supplementary Fig. 6d, e)**. In contrast, both ET-019 and ET-085 significantly increased *SYNGAP1* mRNA expression despite modest effects on productive splicing. ET-085 produced a greater increase in *SYNGAP1* mRNA at lower concentration than ET-019, consistent with our *in vitro* findings **(Supplementary Fig. 6d, e)**.

## Discussion

Here we leverage multiple transcriptome-wide approaches to produce a combined binding and splicing map of an RNA splicing protein in human cortex. Our data identify PTBP binding and splicing of a diverse set of transcripts, while indicating that a propensity of binding is associated with inclusion of AS events to modulate synaptic gene programs. We utilize these PTBP2 splice maps to identify genetic etiologies of neurological disorders that may benefit from splice modulation, and use such a map to guide steric blocking oligonucleotides that disrupt PTBP2-dependent AS of the synaptic gene *SYNGAP1* for potentially therapeutic upregulation.

Transcriptome-wide determination of direct, RBP-mediated AS requires the combination of splicing analysis upon RBP manipulation combined with mapping of RBP binding. We utilized rMATS to quantify AS events across the transcriptome upon depletion of PTBP2 and combined this with CLIP-seq and CLAM-mediated mapping of PTBP2 binding in human cortex and iPSC-neurons. This provides a comprehensive map of PTBP2-binding and AS in human cortical neurons. By examining the positional dependence of PTBP2 binding relevant to AS events, we both confirm previous observations and identify novel features that provide new opportunities for therapeutic intervention for neurodevelopmental disorders. PTBP2 binding within a few hundred nucleotides upstream of a skipped exon is associated with PTBP2-dependent exon silencing but not PTBP2-dependent exon inclusion (**Fig. 3f**). This pattern has been observed for skipped exons previously^12^. One proposed explanation is a competition between PTBP2 and the splicing factor U2AF65^30^, where the presence of PTBP2 could interfere with the use of the cassette exon splice site. Extending this observation, our splicing regulatory maps also show PTBP2 binding upstream of the skipped exon in mutually exclusive splicing events. Another interesting feature to come from our position-dependent splicing analysis is that PTBP2 binding a few hundred nucleotides downstream or upstream of the 5’ and 3’ splice sites, respectively, is associated with intron retention. The conclusion that PBTP2 globally promotes intron retention in cortical neurons extends a previous finding in a mouse neuroblastoma CAD cell line that Ptbp1 represses the splicing of a subset of introns within genes encoding presynaptic proteins^31^. Our analysis in human iPSC-neurons also indicates that PTBP2 generally tends to promote alternative splicing (**Fig. 2a**). Conversely, splicing analysis using splice-sensitive microarrays in Ptbp2 knock-out mouse E18.5 neocortex suggests that Ptbp2 preferentially represses alternative splicing^12^. It is possible that technical differences (i.e. microarray vs. RNA-seq) or cellular context may contribute to the different conclusions.

Our data also indicates a prominent regulation of synaptic gene programs by PTBP2. PTBP2-depletion drives differential expression of 304 genes associated with synaptic organization, function, and plasticity, and we identified concomitant AS and differential gene regulation of several genetic causes of neurological disorders. We focus on *SYNGAP1* as proof of concept for how combined splicing and binding maps can guide ASO-dependent manipulation of an AS event for potential therapeutic gain, for example by preventing NMD of a transcript that drives disease when haploinsufficient. Beyond *SYNGAP1*, **Supplementary Data 2** lists other disease-causing genes that are both alternatively spliced and differentially expressed upon PTBP2 depletion, and which show direct PTBP2 binding near AS events. This should provide a useful roadmap to guide other splice switching strategies for therapeutic manipulation of disease-causing genes.

A limitation of our study is that for differentially spliced Orphanet genes, the identification of the precise AS event and its association with an NMD transcript were generally less clear than with *SYNGAP1*. For example, *GRIN1* (associated with a spectrum of neurodevelopmental disorders) demonstrates PTBP2 binding and alternative splicing proximal to NMD associated events, as well as differential gene expression upon PTBP2 KD. However, the specific exon-exon junction identified as alternatively spliced by rMATS is not annotated in commonly used genome browsers; the more precise identification (and thus targeting) of such events could benefit from long-read sequencing together with short-read sequencing, which was recently demonstrated to help detect many unannotated NMD events across the transcriptome^32^.

Our study also emphasizes the importance for understanding the cell-type and developmental regulation of an AS event for guiding and assessing splice switching therapeutic strategies. NMD events can be tightly developmentally regulated and expressed in a cell-specific fashion. As case in point, *SYNGAP1* expression appears to be strongly repressed by prominent PTBP-dependent AS that induces NMD in both non-neuronal and neuroblastoma cell lines (see **Supplementary Fig. 4a-c**). The NMD inducing AS event becomes increasingly excluded upon neuronal maturation, concomitant with reduced PTBP expression and increased SYNGAP1 levels. By extension, this developmental reduction in AS reduces the potential gain of modulating this event to upregulate SYNGAP1 postnatally. This is evidenced by the >10-fold increase in *SYNGAP1* mRNA via ASO disruption of PTBP binding in HEK293T cells (**Fig. 6e**) compared to far more modest (1.4-fold) upregulation by the same ASO delivered to the postnatal brain (**Supplementary Fig. 6e)**. If *SYNGAP1* haploinsufficiency were identified via genetic testing *in utero*, targeting the high levels of AS during development may have greater potential to restore SYNGAP1 levels (and perhaps improve developmental outcomes) compared to postnatal treatment. Yet such an intervention would require several technical hurdles to be overcome. As *SYNGAP1* is a dosage-sensitive gene (with ∼50% expression likely sufficient to drive pathological phenotypes), even modest postnatal upregulation may still hold therapeutic potential. Our pilot proof-of-concept *in vivo* study motivates further, extensive screening of ASOs to identify oligos with maximal on-target upregulation of *SYNGAP1* and minimal off-target toxicity. As PTBP2 binding sites in non-coding regions are unlikely to be fully conserved between the mouse and human, oligos should target a humanized *SYNGAP1* gene in future *in vivo* studies for optimal translation.

## Methods

### Cell culture and transfection

HEK293T cells were grown in DMEM (Corning #10-013-CV) with 10% Fetal Bovine Serum (Corning #35-010-CV) containing 50 µg/mL gentamicin (Gibco #15750060) and 0.25 µg/mL amphotericin B (Gibco #15290018) at 37 ºC, 5% CO_2_. Cells (2.75 × 10^5^) were seeded in 12-well plates one day before transfection, which was performed with 50, 100 or 200 nM of ASO using Lipofectamine RNAiMax reagent (Invitrogen #13778100) or Lipofectamine 2000 (Invitrogen #11668027) according to manufacturer’s instructions. For decoy experiments, HEK293T cells were transfected with 5 µM of the specified decoy. Total RNA was isolated from HEK293T cells 24h or 48h after transfection. Total protein was extracted 48h post-transfection.

PTBP1 and PTBP2 knockdown experiments in HEK293T cells were performed with TriFECTa kit DsiRNA Duplex (IDT) using hs.Ri.PTBP1.13 (named siPTBP1) and hs.Ri.PTBP2.13 (named siPTBP2) predesigned siRNAs, respectively. Predesigned NC-1 siRNA (IDT #51-01-14-04) was used as negative control (named siSc). siRNAs were transfected with Lipofectamine RNAiMax reagent using the concentrations recommended by the manufacturer and total RNA and protein were isolated 48h post-transfection.

Ptbp1 was knocked down in mouse Neuro2A (N2a) cells using LNA gapmer ASOs. Briefly, cells were seeded at 75,000 cells per well in a 24-well plate and transfected the following day with 3 µL of 20-µM Ptbp1 or control ASO with 0.6 µL of P3000 reagent and 1.5 µL of Lipofectamine 3000 reagent (Invitrogen) per well, according to the manufacturer’s instructions. After 48 hours, RNA was extracted using 1 mL of TRIzol (Invitrogen) according to manufacturer’s instructions.

SH-SY5Y cells were grown in 1:1 Ham’s F12:EMEM (Gibco #11765-047 and ATCC #30-2003, respectively) with 10% Fetal Bovine Serum (Corning #35-010-CV) containing 50 µg/mL gentamicin (Gibco #15750060) and 0.25 µg/mL amphotericin B (Gibco #15290018) at 37 ºC, 5% CO_2_. ASO (20 µM) was delivered into cells (1 × 10^6^) by electroporation in a Nucleofector device (Lonza) using the Amaxa Cell Line Nucleofector Kit V (Lonza #VCA-1003) and following the recommended instructions for SH-SY5Y nucleofection. Total RNA was isolated from SH-SY5Y 24 h after nucleofection.

### Generation of iPSC-derived neurons

For generation of mature iPSC-neurons, we used a protocol, modified from a combination of previous studies^33,34^, described in further detail, below. The CHOP-WT10 cell line was maintained as an iPSC culture^35^ and subsequently transitioned to feeder-free cultures and maintained with mTeSR1 (StemCell Technologies) on hESC qualified matrigel (Corning). Feeder-free iPSC stocks were cryopreserved in 90% FBS/ 10% DMSO at a minimum of 2 passages after feeder-free transition. Feeder-free stocks of CHOP-WT10 were passaged in mTeSR1 prior to the initiation of differentiation.

Prior to differentiation, plates were coated with growth factor reduced (GFR) Matrigel (Corning), where 1 mg of GFR Matrigel was added to 24 ml of DMEM/F12 (Invitrogen) and 2 ml of the mixture was added per 35 mm well and incubated at 37 ºC for a minimum of 1 h^36^. iPSCs were passaged with mTeSR1 onto the GFR Matrigel-coated plates at a density of approximately 50,000 cells/cm^2^ well. When cultures were a minimum of 60% confluent, differentiation of iPSCs into neural progenitor cells (NPCs) was initiated (indicated as “Day 0”), as previously described in^33^ with modification. Briefly, cultures were treated with daily media changes containing SB431542 (10 µM; Tocris), LDN193189 (1 µM, Stemgent), IWR1 (1.5 µM; Tocris) and supplemented with B27 without vitamin A (Invitrogen) and passaged at days 4 and 8 of differentiation. Cells were passaged with accutase (Invitrogen) and replated on to GFR Matrigel-coated plates in the appropriate media, containing Y27632 (10 µM), at the approximately densities: 210,000/cm^2^ (Day 4); 155,000/cm^2^ (Day 8). After 8 days of daily treatment, forebrain neural ectoderm was confirmed by expression of FOXG1, SOX2, and PAX6 by flow cytometry (data not shown).

From Day 8 to 14 of differentiation, NPCs were expanded in Invitrogen Neural Expansion Media, containing Neural Induction Supplement in 50% Advanced DMEM/F12 and 50% Neurobasal Medium [Invitrogen, as per manufacturer instructions;^34^]. NPC identity was confirmed at Day 14 by expression of FORSE-1 and rosette morphology (data not shown) and NPCs were cryopreserved in 90% FBS/ 10% DMSO for future use. NPCs continued directly or were thawed from cryopreserved NPCs for subsequent differentiation steps. NPCs were plated on to GFR Matrigel coated plates at 285,000/cm^2^ into N2B27(−): 2 parts DMEM/F12 (Invitrogen), 1 part Neurobasal Medium (Invitrogen) containing 1/3X N2 supplement (Invitrogen), 2/3X B27 without vitamin A (Invitrogen), 1X glutamine (Invitrogen), 50 µM beta-mercaptoethanol (Invitrogen) and 25 ng/ml of Activin A (Bio-Techne), followed by daily media changes of the same media with supplements. After 8-9 days of N2B27(−) + Activin A, the majority of cells appeared as terminally differentiating neurons, and were therefore passaged with 0.5 mM EDTA, and replated at a density of approximately 155,000/cm^2^ on to plates coated with poly-D-lysine (10 µg/mL, Sigma), followed by GFR matrigel, as above, into N2B27(+): 2 parts DMEM/F12 (Invitrogen), 1 part Neurobasal Medium (Invitrogen) containing 1/3X N2 supplement (Invitrogen), 2/3X B27 with vitamin A (Invitrogen), 1X glutamine (Invitrogen), 50 µM beta-mercaptoethanol (Invitrogen), brain-derived neurotrophic factor (20 ng/mL) and glial-derived neurotrophic factor (20 ng/mL). Half-media changes were performed with the same media and supplements 3 times a week until final harvest. Neuronal identity was confirmed by immunofluorescence staining of MAP2 (Sigma) and Tuj1 (Biolegend) (**Supplementary Fig. 1a**). qPCR analyses confirmed expression of TBR2, TBR1, CTIP2, SATB2, VGLUT1, and VGLUT2 in the final neuron population (**Supplementary Fig. 1b**). Experiments were performed when neurons matured to day 40-50 of differentiation, or the equivalent, in cases where neurons were cryopreserved at day 14.

For Western blot analysis, cells were harvested in 1.5X Laemmli sample buffer [75 mM Tris-HCl, pH 7.5 (Invitrogen #15567027), 3% SDS (Sigma #L5750), 15% glycerol (Amresco #M152), 3.75 mM EDTA, pH 7.4 (Bio-Rad #1610729)]. Cells were harvested in RNA lysis buffer (Zymo Research #R1055) for RT-PCR and qPCR analysis or TRIzol (Invitrogen #15596-026) for RNA-seq.

### Cycloheximide and ASO treatment

To evaluate SYNGAP1 as a target for NMD, we incubated iPSC-neurons with 50 µM of cycloheximide (CHX) (Sigma-Aldrich #01810) for 3h. ASO treatments in mature iPSC-neurons were performed by gymnotic delivery of the ASO for 7 days, maintaining scheduled half-media changes until harvest.

ASO and decoy oligonucleotides were purchased from IDT. Gapmer ASOs were obtained from Qiagen. Sequences and chemistry information for all oligonucleotides can be found in **Supplementary Data 3**.

### RNA-seq library preparation and sequencing

PTBP2 gapmer-treated (5 µM PTBP2_7), control gapmer-treated (5 µM NegA), or untreated iPSC-derived neurons were lysed with TRIzol Reagent (Invitrogen 15596-026). Chloroform was added to the lysate, and samples were centrifuged at 4°C to separate phases. After addition of ethanol, the aqueous phase was applied to an RNeasy Mini column (Qiagen #74104). On-column DNAse treatment was performed using Qiagen DNAse Set (Qiagen #79254) at room temperature for 15 minutes. Purified RNA was analyzed on Agilent TapeStation systems, using D1000 Screen Tape & Reagents, and then quantified with a Qubit BR kit (Thermo Fisher #Q10210). Poly(A)+ RNA transcript was isolated from 1 ug purified total RNA (RNA integrity number >9.0) with NEBNext poly(A) mRNA magnetic isolation module (New England Biolabs #E7490). RNA-seq libraries were prepared with NEBNext Ultra Directional RNA library preparation kit for Illumina (New England Biolabs #E7420S) according to the manufacturer’s instruction. The samples were sequenced using NovaSeq 6000 SP Reagent Kit v1.5 (300 cycles) with 150-bp paired-end reads at the Sequencing Core at Children’s Hospital of Philadelphia.

### RNA-seq data processing for differential gene expression

#### Genome mapping

RNA-seq libraries were demultiplexed, and adapter sequences were removed with Cutadapt v1.18. Sequence quality was assessed by FastQC v0.11.2. Transcriptome indices were prepared for Salmon v1.5.2 using a decoy-aware transcriptome file (gencode.v38.transcripts.fa with GRCh38.primary_assembly.genome.fa genome as decoy). Transcripts were quantified from paired-end reads in mapping-based mode (selective alignment, --libType ISR --gcBias -- validateMappings).

#### Differential gene expression

Salmon transcript counts were imported into R and aggregated to the gene level using the tximeta^37^ and SummarizedExperiment^38^ packages in Bioconductor. Differential expression analysis was performed using DESeq2^39^ with Untreated or NegA as the reference condition, as indicated. The p-values attained by the Wald test were corrected for multiple testing using the Benjamini-Hochberg method. An adjusted pval cutoff of 0.05 was used for significance. Log2 fold change shrinkage was applied using the apeglm algorithm^40^. Ensembl gene annotations were added. Overrepresentation analysis was performed using the clusterProfiler package in Bioconductor^41^. All genes evaluated for differential expression were used as the background dataset for testing. Overrepresentation analysis specifically of synaptic genes was performed using SynGO annotated genes^19^ release 1.1 with brain-expressed genes as the background set.

### RNA-seq data processing for alternative splicing analysis

#### Genome alignment

Demultiplexed, paired-end reads were aligned to the human genome (GRCh38.primary_assembly.genome.fa) with STAR v2.7.1a in two-pass mode^42^. For first-pass alignment, GENCODE basic gene annotation (gencode.v38.basic.annotation.gtf) was used (−-outSAMstrandField intronMotif --outFilterType BySJout --alignSJoverhangMin 8 -- alignSJDBoverhangMin 3 --outFilterMismatchNoverReadLmax 0.04 --alignIntronMin 20 -- alignIntronMax 1000000 --alignMatesGapMax 1000000 --scoreGenomicLengthLog2scale 0). The output splice junction files were concatenated and filtered (to remove junctions on chrM, non-canonical junctions, junctions supported by multi-mappers or by too few reads) then used in on-the-fly second-pass alignment (−-outSAMstrandField intronMotif --outSAMattributes NH HI AS NM MD --outFilterType BySJout --alignSJoverhangMin 8 --alignSJDBoverhangMin 3 -- outFilterMismatchNoverReadLmax 0.04 --alignIntronMin 20 --alignIntronMax 1000000 -- alignMatesGapMax 1000000 --scoreGenomicLengthLog2scale 0 --quantMode TranscriptomeSAM GeneCounts). Alignments were filtered with samtools v0.1.19 to remove unmapped reads and reads mapping to 10 or more locations. Alignment quality was assessed with Qualimap v2.2.2-dev.

#### Splicing analysis

Differential splicing analysis was performed using rMATS v4.1.1 with bam files as input (group 1 Untreated, group 2 PTBP2 KD). Prep and post steps were run (−-gtf gencode.v38.primary_assembly.annotation.gtf -t paired --libType fr-firststrand --readLength 151 --variable-read-length --novelSS --allow-clipping) followed by the statistical model (−-cstat 0.01). rMATS corrects for multiple testing using the Benjamini–Hochberg FDR method. rMATS output files were then loaded into R with the maser package in Bioconductor^43^. Downstream analysis was performed using the junction counts only file, and splicing events were filtered by coverage (average read counts of 20 or more).

#### Annotation of disease relevance

A list of genes associated with rare diseases were downloaded from the Orphadata repository of Orphanet (http://www.orphadata.org).

#### Visualization

Bigwig files were prepared using Yeo lab’s makebigwigfiles [https://github.com/YeoLab/makebigwigfiles]. Visualization was performed in R using the Gviz package in Bioconductor^44^.

### Human cortex RNA-seq data

The Genotype-Tissue Expression (GTEx) Project was supported by the Common Fund of the Office of the Director of the National Institutes of Health, and by NCI, NHGRI, NHLBI, NIDA, NIMH, and NINDS. The data used for the analyses described in this manuscript were obtained from: Brain Front Cortex (gtexCovBrainFrontalCortexBA9) table from the GTEx RNA-seq Coverage track on the UCSC genome browser on 5/6/2022.

### eCLIP for iPSC-neurons and postmortem human brain

Postmortem human brain samples (frontal cortex, Brodmann area 9) were obtained from Johns Hopkins University (Baltimore, MD). eCLIP was performed as described previously ^45,46^ except the adaptor sequences were from^47^. For iPSC-derived neurons, 3x 6-well plates were used for 3 eCLIP replicates. The differentiated neurons seeded in 1x 6-well plate (∼1×10^6^ cells/well) were irradiated with ultraviolet (UV) light once at 400mJ/cm^2^ and once at 200 mJ/cm^2^ on iced water. For postmortem human brain BA9 region, the tissue from each of three different donors was first pulverized in liquid nitrogen, and the powder in a chilled 10-cm tissue culture plate on dry ice was UV-crosslinked at 400 mJ/cm^2^ three times^46^. Crosslinked neurons or brain tissues were washed 1x with pre-chilled DPBS (Corning #21-031-CV). Pellets were frozen in -80 °C or directly lysed in eCLIP lysis buffer (50 mM Tris-HCl pH 7.4, 100 mM NaCl, 1% NP-40 (Igepal CA630), 0.1% SDS, 0.5% sodium deoxycholate) containing proteinase inhibitor and RNAse inhibitor. Cell or tissue lysis was assisted with sonication by using a Bioruptor on the low setting for 5 min, cycling 30 s on and 30 s off. For each 1 mL lysate, 5 µL Turbo DNAse I (Thermo Fisher #AM2239) and high dose RNAse I (Thermo Fisher #AM2295; 1:5 diluted in cold DPBS) or low dose RNAse I (1:50 dilution for neurons, and 1:100 dilution for brain) were added. The microspin tubes were placed in a Thermomixer preheated to 37 °C for exactly 5 min, shaken at 1,200 rpm, and then put on ice to terminate the reaction. Cell lysates were spun for 20 min at full speed in a pre-chilled centrifuge.

The cleared lysate was mixed with Dynabeads Protein G (Thermo Fisher #10003D) pre-conjugated with 10 µg PTBP2 antibody (EMD Millipore #ABE431) and incubated in a 4 °C cold room overnight. Immunoprecipitated-RNA was dephosphorylated with FastAP enzyme (Thermo Scientific #EF0652) and T4 PNK (New England BioLabs #M0201L), and an adapter labeled with an IRDye®800CW fluorochrome (/5Phos/rNrNrNrN rArGrA rUrCrG rGrArA rGrArG rCrArC rArCrG rUrCrU rGrArA rArA/3IR800CWN/) was ligated to the 3′ end. Labeled RBP-RNA complexes were eluted using 1x LDS loading buffer (Thermo Fisher) and resolved on 4-12% Bis-Tris gels, transferred to nitrocellulose membrane, and imaged. RNA-protein complexes were visualized with LI-COR Odyssey imaging system, and a region including ∼75 kDa above protein size (by comparing with high RNAse I sample) was cut from the membrane. RNA was isolated from the membrane via protease K/SDS treatment. After reverse transcription with a modified primer (5’-TTC AGA CGT GTG CTC TTC CG-3’), a 5′ adapter containing 8-nt UMI (/5phos/NNNNNNNN AGATCGGAAGAGCGTCGTGTAGGG/3ddC/) was ligated to cDNA. All of the remaining steps were essentially performed according to the eCLIP procedure^45^. The uniquely barcoded eCLIP samples were prepared and analyzed on Agilent Tapestation D1000. The final libraries were pooled and sequenced using NovaSeq 6000 SP Reagent Kit v1.5 (300 cycles) with 150-bp paired-end reads at the Sequencing Core at Children’s Hospital of Philadelphia.

### eCLIP-seq data processing

#### Genome alignment

eCLIP libraries were demultiplexed, and inline random 8-mers were removed from the start of read 1 and appended to the read name using a script modified from the Yeo lab’s eclipdemux [https://github.com/YeoLab/eclipdemux]. Inline random 4-mers were removed from the start of read 2 using Trimmomatic v0.32. Adapter sequences were removed in two rounds with Cutadapt v1.18 using commands recommended by ENCODE’s eCLIP-seq Processing Pipeline. Sequence quality was assessed by FastQC v0.11.2. Paired-end reads were aligned to the human genome (GRCh38.primary_assembly.genome.fa, gencode.v39.primary_assembly.annotation.gtf) with STAR v2.7.1a (−-alignEndsProtrude 15 ConcordantPair --outFilterMultimapNmax 100 --outSAMattributes NH HI AS NM MD). Multimappers were further filtered prior to peak calling (see below). rRNA and tRNA tracks were downloaded from UCSC Table Browser (RepeatMasker’s rmsk track and filtering for rRNA or tRNA), and rRNAs and tRNAs were removed in a two-step process. First, bedtools intersect (v2.15.0, -f 0.90) was used to identify all rRNA/tRNA reads. In the second step, qnames from round 1 were used to mask potentially multi-mapping rRNAs/tRNAs in the alignment file using Picard Tools v1.141 FilterSamReads.

#### Peak calling

PCR duplicate removal was performed using collapse_duplicates.py from the Xing lab^21^ with modifications. Peak calling was performed using CLAM v1.2^21^. Briefly, CLAM preprocessor was used to separate multi-mapped and uniquely mapped reads. Multi-mapped reads (−-max-multihits of 10) were re-aligned to the genome with a probability. CLAM peakcaller was used in multi-replicate mode with size-matched input as background (n = 3 replicates each for IP and input) using default values for --qval-cutoff (0.05), --fold-change (2-inf), and --binsize (50). CLAM corrects for multiple testing using the Benjamini–Hochberg FDR method. Finally, peak annotation was performed with CLAM peak_annotator.

#### Peak analyses

De novo motif finding was performed using Homer v4.11 (−len 5,6,7 -size 100 -S 10). Peak distribution by genomic feature was calculated using RSeQC v4.0.0 read_distribution.py. Overrepresentation analysis was performed using the clusterProfiler package in Bioconductor^41^. All genes evaluated for differential expression were combined with genes with CLIP-seq peak calls and used as the background dataset for testing. Overrepresentation analysis specifically of synaptic genes was performed using SynGO annotated genes^19^ release 1.1 with brain-expressed genes as the background set.

#### Splicing regulatory maps

Splicing regulatory maps were generated using RBPmaps^23^ (−-normalization_level 0, --sigtest zscore) with CLAM peaks and significant AS events identified by rMATS (pval < 0.05, FDR < 0.1, IncLevelDifference >= |0.05|). The rMATS output was first subsetted by overlaps to eliminate overlapping events using a script provided by RBPmaps (subset_rmats_junctioncountonly.py). Background controls provided by RBPmaps were used for SE (HepG2_native_cassette_exons_all), A5SS (HepG2-all-native-a5ss-events), and A3SS (HepG2-all-native-a3ss-events) events.

### Single bolus ICV injection in neonate mice

C57BL/6J male and female mice were used in this study. All mice were maintained on a 12:12-h light:dark cycle and mothers had *ad libitum* access to food and water throughout the experiments. Lyophilized ASO was reconstituted in 1x PBS (Thermo Fisher #10010023) and diluted to the desired concentration. For ICV injection in P2 mice, pups were cryoanesthetized and then placed on a cold metal plate. A 30-gauge blunt needle mounted on a 10 µL Hamilton syringe was used to pierce the skull one-third the distance between lambda and the eye and at a depth of 2 mm. 2 μL of ASO or PBS was injected slowly into the right cerebral lateral ventricle. Injected mice were rewarmed and quickly returned to the nest and observed daily for survival and signs of stress. Animals were sacrificed at P7, and brain sectioned into R and L hemispheres after discarding the olfactory bulbs, cerebellum, and brainstem. Tissue was flash-frozen in liquid nitrogen and stored at -80ºC until further processing. The right hemisphere was used for RNA analysis.

Animal care and use procedures were approved and performed in accordance with the standards set forth by the Children’s Hospital of Philadelphia Institutional Animal Care and Use Committee and the Guide for the Care and Use of Laboratory Animals published by the US National Institutes of Health.

### RNA extraction and cDNA synthesis for RT-PCR and qPCR

Total RNA from cells was isolated using Quick-RNA Miniprep Kit (Zymo Research #R1055) following manufacturer’s instructions, including DNase treatment step. RNA was eluted in a final volume of 50 µL of RNase-free water.

Total RNA from mouse brain tissue was extracted using TRIzol. Tissue section (right hemisphere) was mixed with 1 mL of TRIzol reagent (Invitrogen #15596018) and a 5 mm stainless steel bead (Qiagen #69989) in an RNase-free microcentrifuge tube. Brain tissue was then homogenized in a TissueLyser LT homogenizer (Qiagen #85600) for 5 min at 50 Hz. The homogenate was centrifuge at 12,000 x g for 5 min at 4 °C and then seated for an additional 5 min to precipitate insoluble debris. The supernatant was transferred to a new tube, mixed with 200 µL of chloroform (Acros Organics #190764), shaken vigorously and centrifuged at 12,000 x g for 15 min at 4 °C. Aqueous phase was transferred to a new tube containing 500 µL of ice-cold isopropanol (Sigma-Aldrich #190764) followed by incubation for 10 min on ice and centrifugation at 12,000 x g for 10 min at 4 °C to precipitate RNA. Supernatant was discarded and 1 mL of ice-cold 75% ethanol (Decon laboratories #2701) was added to wash the pellet followed by centrifugation at 7,500 x g for 5 min at 4 °C. Supernatant was again discarded and RNA pellet was air-dried for 20 min. RNA was resuspended in 200 µL of RNase-free water and allowed to reconstitute for 10 min at 56 °C.

RNA concentration was determined by measuring OD_260_ nm absorbance in a Synergy HTX reader (Biotek). cDNA synthesis was performed using the SuperScript IV First-Strand Synthesis System with ezDNase Enzyme (ThermoScientific #18091300) using random hexamer primers according to manufacturer’s instructions. The ezDNase treatment step was performed for all conditions.

### RT-PCR

PCR reaction was prepared in a 0.2-mL tube by mixing the following reagents: 5 µL cDNA template, 1x MyTaq reaction mix (BIO-LINE #25042), forward and reverse primers (0.4 µM each) and nuclease-free water in a final volume of 25 µL.

For SYNGAP1 (exon 10-11), PCR analysis to assess both the productive and non-productive transcript was performed using forward primer 5’-AATTCATCCGTGCTCTGTATGA-3’ and reverse primer 5’-AAGAGACTGGGCGACATAATC-3’. PCR cycling conditions were 15 s at 95 ºC for denaturation, 15 s at 64 ºC for annealing, and 30 s at 72 ºC for extension for 26 cycles. The predicted molecular weights of the PCR products were 289 bp for the productive transcript (containing exon 10 and 11) and 465 bp for the non-productive transcript (containing exon 10, 11x and 11).

For SYNGAP (exon 14), PCR analysis to assess inclusion of exon 14 was performed using forward primer 5’-TCCTGAAGCTGGGTCCACTG-3’ and reverse primer 5’-GGGTGGCTTTTCCTTGGTTG-3’. PCR cycling conditions were 15 s at 95 ºC for denaturation, 15 s at 68 ºC for annealing, and 30 s at 72 ºC for extension for 26 cycles. The predicted molecular weights of the PCR products were 221 bp for exon exclusion (containing exon 13 and 15) and 263 bp for the exon inclusion (containing exon 13, 14 and 15).

For SYNGAP (exon 18-19), PCR analysis to assess both the productive and non-productive transcript was performed using forward primer 5’-AGGCAGAGAAGGATTCCCAGA-3’ and reverse primer 5’-TCACACGCGGGTTTGTTGG-3’. PCR cycling conditions were 15 s at 95 ºC for denaturation, 15 s at 64 ºC for annealing, and 30 s at 72 ºC for extension for 26 cycles. The predicted molecular weights of the PCR products were 174 bp for the productive transcript (containing exon 17, 18 and 19) and 254 bp for the non-productive transcript (containing exon 17, 18, 19x and 19).

For ms-Syngap1 (ex10-11), PCR analysis to assess both the productive and non-productive transcript was performed using forward primer 5’-AGACCCCATCAAGTGCACAG-3’ and reverse primer 5’-GCCTGTCAGCAATGTCCTCT-3’. PCR cycling conditions were PCR cycling conditions were 15 s at 95 ºC for denaturation, 15 s at 65 ºC for annealing, and 45 s at 72 ºC for extension for 25 cycles. The predicted molecular weights of the PCR products were 185 bp for the productive transcript (containing exon 10 and 11) and 356 bp for the non-productive transcript (containing exon 10, 11x and 11).

For ms-Scn1a, PCR analysis to assess both the productive and non-productive transcript was performed using forward primer 5’-CAGTTTAACAGCAAATGCCTTGGGTT-3’, reverse primer 5’-AAGTACAAATACATGTACAGGCTTTCCTCATACTTA-3’ and cycling conditions from^6^. The predicted molecular weights of the PCR products were 498 bp for the productive transcript (containing exon 21, 22, 23 and 24) and 562 bp for the non-productive transcript (containing exon 21, 21x, 22, 23 and 24).

PCR products were mixed with 1x GelRed prestain loading buffer (Biotium #41010), separated on a 2% or 4% agarose gel (Seakem LE agarose, Lonza #50004) by electrophoresis (120 V, up to 1.5 h) and imaged using an AlphaImager system (AlphaInnotech). Gels were quantified with Image Studio Lite software (LI-COR). Productive splicing (%) was quantified as the background-subtracted integrated intensity of the productive band divided by the sum of the background-subtracted integrated intensities of the productive + nonproductive band x 100. Percent spliced in (PSI) was quantified as the background-subtracted integrated intensity of the band containing the element of interest divided by the sum of the background-subtracted integrated intensities of the band containing + band lacking the element of interest x 100.

### qPCR

Probe-based qPCR was prepared by mixing the following reagents: 1 µL of cDNA, 1X PrimeTime Gene Expression Master Mix (IDT #1055772), 1X primers/probe mix and nuclease-free water to a final volume of 10 µL. Three technical replicates were performed for each sample. qPCR was carried out on a QuantStudio 3 or QuantStudio 5 Real-Time PCR System (ThermoFisher) with a passive reference of ROX using the following cycling conditions: 95 ºC for 3 min for 1 cycle, 95 ºC for 5 s and 60 ºC for 30 s for 40 cycles. ΔCt was calculated by subtracting the average Ct of the reference gene from the average Ct of the gene of interest for each sample. ΔΔCt values were obtained by subtracting the average ΔCt value of control samples from the ΔCt of the test samples, and then converted into 2^-ΔΔCt^ to obtain the fold change of gene expression.

qPCR data were normalized to ATP5F1 (**Fig. 1c, 4d, 4e**), GAPDH (**Fig. 5b**), RPL4 (**Fig. 5c, 5g, 6c, 6e, Supplementary Fig. 1d, Supplementary Fig. 5a, b, c and e**) or mouse Atp5f1 (**Supplementary Fig. 6c, e**).

The following qPCR probes were used for mRNA expression analyses: SYNGAP1 exon 10-11 (Hs00405348_m1, ThermoFisher), SYNGAP1 exon 16-17 (Hs.PT.58.4622325, IDT, used for CHX experiments only), RPL4 (Hs00973293_g1, ThermoFisher), ATP5F1 (Hs01076982_g1, ThermoFisher), GAPDH (Hs00266705_g1, ThermoFisher), PTBP1 (Hs.PT.58.25863276, IDT), PTBP2 (Hs.PT.58.20884110, IDT), ms-Syngap1 (Mm01306145_m1, ThermoFisher), ms-Scn1a (Mm00450583_mH, ThermoFisher), ms-Atp5f1 (Mm05814774_g1, ThermoFisher).

### qPCR of neuronal markers

iPSC-neurons were harvested between day 40-50 of differentiation, or equivalent, in RNA lysis buffer (Zymo Research) and RNA was purified with the Quick-RNA Miniprep kit (Zymo Research R1054), as per manufacturer’s instructions. Additional gDNA was subsequently removed with the TURBO DNase free kit (ThermoFisher), as per manufacturer’s instructions. cDNA was generated with a mix containing 9 µL RNA, 1 µL N6 random hexamers (Invitrogen) that were held at 70 ºC for 5 min, followed by 4 ºC. 4 µL of 5X First-Strand Buffer, 1 µL 0.1M DTT, 1 µL 10mM dNTPs mix, 1 µL RNaseOUT, 0.75 µL superscript III (each from Invitrogen), and 2.25 µL RNase/DNase-free water were added and the mixture was subjected to five-stepwise incremental temperature increases from 25 ºC to 70 ºC. The resulting reaction was kept at 4 ºC until use and diluted 1:10 in RNase/DNase-free water.

qPCR was performed with 1 µL of a primer mix (2 µM each of forward and reverse primer), 2.5 µL SYBR green (ThermoFisher), and 1.5 µL of cDNA. Mastermixes of the primer mix + SYBR green were added to all applicable wells prior to the addition of cDNA, which was added individually in triplicate. A standard curve of gDNA (10, 1, 0.1 ng/mL) generated from H9 embryonic stem cells was also examined against each primer pair. All primer pairs were generated to be within exon such that they could be used to calculate a concentration of cDNA transcript in the given sample. Each transcript concentration was standardized against the concentration of GAPDH in the sample. Note that GAPDH primer sets additionally bind to 3 pseudo-genes, resulting in additional gDNA binding and thereby allowing for assessment of GAPDH in the same linear range as other transcripts.

qPCR samples were analyzed on a QuantStudio 5 System (Applied Biosystems) with a passive reference of ROX and analyzed by QuantStudio 5 software using the Relative Standard Curve setting.

Primer sequences used for assessing expression of neuronal markers can be found in **Supplementary Data 4**.

### Immunofluorescence

iPSC-neurons were plated on to German-glass coverslips (Electron Microscopy services) for the terminal differentiation stage and then fixed at Day 46 of differentiation with 4% paraformaldehyde for 30 min at room temperature. Cells were permeabilized with PBS + 0.3% Triton X-100 (Sigma) for 10 min. Cells were then blocked with Animal-Free Blocking solution (Cell Signaling Technology) for 1h at room temperature, followed by primary antibody diluted in PBS + 3% BSA (Sigma A1470) overnight at 4 ºC. Primary antibodies included: MAP2 (Sigma M1406, 1:500), Tuj1 (Biolegend 801201, 1:500), PSD-95 (Cell Signaling 3450, 1:200). Cells were subsequently treated with secondary antibodies goat anti-IgG1-Alexa488 (for MAP2), goat anti-IgG2A-Alexa647 (for Tuj1), goat anti-rabbit Alexa568 (for PSD-95) (each 1:500, Invitrogen) diluted in PBS + 3% BSA for 2h at room temp, follow by post-fixation with 4% paraformaldehyde for 15 min at room temp. Cells were counterstained with Hoechst-33342 (Invitrogen, 1:2000) in PBS for 15 min at room temperature, and mounted with Fluoromount-G (Southern Biotech). Washes with PBS were performed between each step, and a final wash with water was performed prior to mounting. Images were taken on a Leica DMi4000 inverted microscope, outfitted with a PL APO 40X objective and Leica DFC340 FX camera.

### Calcium transient measurement in iPSC-neurons

iPSC-neurons were cultured on Mattek (#P35G-1.5-20-C) dishes. Prior to imaging, medium was replaced with BrainPhys neuronal medium^48^. iPSC-neurons were incubated with 3 µM Fluo-4 (Life Technologies #F14201) at 37°C for 20 min and imaged directly following washout on an LSM Zeiss 880 inverted confocal microscope using a 63×oil 1.4 numerical aperture objective and a 488-nm argon ion laser. Calcium transients were measured at rest and following electrical stimulation with 20 Hz trains of depolarizing field stimuli lasting 10 s, with 20 s of rest between trains (IonOptix).

### Western blot

Total protein extraction from cells was performed using 1.5x Laemmli buffer [15% glycerol (Amresco #M152), 3% SDS (Sigma #L5750), 3.75 mM EDTA (Bio-Rad #1610729) and 75 mM Tris, pH 7.5 (Invitrogen #15567027)] followed by 10 min incubation at 95 ºC. Protein extracts were quantified with Pierce BCA protein assay kit (ThermoFisher #23227) according to manufacturer’s instructions, diluted to the same final concentration, mixed with 1x Orange G dye (Sigma #O3756) containing 10% β-mercaptoethanol (Sigma #M3148) and incubated 10 min at 100 ºC before loading. Precast 4-15% TGX protein gels (Bio-Rad) were loaded with 10-30 µg of total protein lysate and run for 1h at 110-135V. Transfer was performed with a Trans-Blot Turbo Transfer system (Bio-Rad) using the pre-determined high molecular weight transfer protocol for SYNGAP1 (10 min, 2.5 A constant) or the pre-determined mixed molecular weight transfer protocol for PTBP2 (7 min, 2.5 A constant).

For SYNGAP1 blots, proteins were transferred to a 0.45 µm low fluorescence PVDF membrane (Bio-Rad #1704275) and blocking was performed using 5% ECL prime blocking reagent (Cytiva #RPN418) for at least 1h at room temperature. Incubation with primary antibodies (diluted in blocking buffer) was carried out overnight at 4 ºC. Rabbit anti-SYNGAP1 (Cell Signaling Technology #5539S, 1:1000 dilution), previously validated^6^ and mouse anti-ATP5F1 (Abcam #ab117991, 1:1000 dilution or Santa Cruz Biotechnology #sc-514419, 1:500 dilution) were used. Membrane was then rinsed with 1x TBST (diluted from TBST-10X, CST 9997) 4 times for 5 min. Incubation with secondary antibodies (diluted in blocking buffer) was performed at room temperature for 1h. Anti-rabbit-HRP (Cell Signaling Technology #7074S, 1:5000 dilution) and anti-mouse-HRP (Cell Signaling Technology #7076S, 1:5000 dilution) were used. Membrane was rinsed again with 1x TBST 4 times for 5 min before imaging. Blots were developed using SuperSignal West Femto Maximum Sensitivity Substrate (ThermoScientific #34095) and detection was carried out on a GBox imaging system (Syngene). Blot images were exported in 16-bit grayscale format for further analysis.

For PTBP2 blots, proteins were transferred to a 0.2 µm nitrocellulose membrane (Bio-Rad #1704271) and blocking was performed using Intercept (TBS) Blocking Buffer (LI-COR #927-60001) for at least 1h at room temperature. Incubation with primary antibodies (diluted in blocking buffer containing 0.1% Tween-20) was carried out overnight at 4 ºC. Rabbit anti-PTBP2 (EMD Millipore #ABE431, 0.5 µg/mL), mouse anti-GAPDH (D4C6R) (Cell Signaling Technology #97166S, 1:1000 dilution) and mouse anti-ATP5F1 (as above) were used. Membrane was then rinsed with 1x TBST 4 times for 5 min. Incubation with secondary antibodies (diluted in blocking buffer containing 0.1% Tween-20) was performed at room temperature for 1h. IRDye 680RD anti-rabbit (LI-COR #926-68073, 1:10,000 dilution) and IRDye 800CW anti-mouse (LI-COR #926-32212, 1:10,000 dilution) were used. Membrane was rinsed again with 1x TBST 4 times for 5 min and imaged on Odyssey Imager (LI-COR) using a resolution of 169 µm.

Western blot quantifications were normalized to ATP5F1 or GAPDH according to LI-COR’s Housekeeping Protein Normalization Protocol.

### Statistical analyses

Measurements were taken from distinct samples. The number of samples is stated explicitly in the figure legend and represented as individual points for bar graphs. Statistical significance for two-tailed Student’s t-test or one-way analysis of variance (ANOVA) with Tukey’s or Dunnett’s post-test was defined as p < 0.05 (or adjusted p value of <0.05 or FDR of <0.05 for sequencing data) and calculated using GraphPad Prism 9.3 software, R, or software used for sequencing data. Data plots were prepared using GraphPad Prism 9.3 software and R. Individual statistical tests applied to each data set are given in the respective figure legends or methods section. Bar graphs represented as mean values ± SEM.

## Supporting information

Description of Supplementary Files

## Data availability

The datasets generated during the current study are available in the GEO repository with accession codes GSE206661, GSE206650, and GSE206660.

## Acknowledgements

We thank S. Zhang, M. Reilly and J. Petrosino (all from University of Pennsylvania) for providing feedback on the manuscript. We thank D. Leib (Children’s Hospital of Philadelphia) for providing a reagent.

## Author Contributions

J.M.D-M., A.J.F., E.A.W., C.C., D.A.A., A.B., B.L.D., and B.L.P. contributed to experimental design. J.M.D-M., A.J.F., E.A.W., C.C., D.A.A., A.B., and L.V.D. performed the experiments and collected the data. E.A.W. and L.V.D. maintained, differentiated, and treated iPSCs and iPSC-neurons. C.C. performed CLIP-seq and prepared libraries for sequencing. D.A.A. performed *in* vivo injections of ASOs. J.M.D-M., A.J.F., E.A.W, and A.B. contributed to data analysis. J.M.D-M. performed bioinformatics analysis with feedback from P.T.R.. D.L.F., B.L.D., and B.L.P. supervised the project. J.D.M, A.J.F, and B.L.P. prepared the manuscript with input from E.A.W, C.C., D.A.A., E.A.H., D.L.F., and B.L.D. All authors discussed the results and participated in the critical review and revision of the manuscript.

## Materials & Correspondence

Correspondence and material requests should be addressed to B.L.P. and B.L.D..

## Intellectual Property

Provisional patent 22-9943 has been filed on behalf of J.M.D-M., A.J.F., B.L.D., B.L.P..

