## Supplementary Figures for "Mapping PTBP splicing in human brain identifies targets for therapeutic splice switching including *SYNGAP1*"

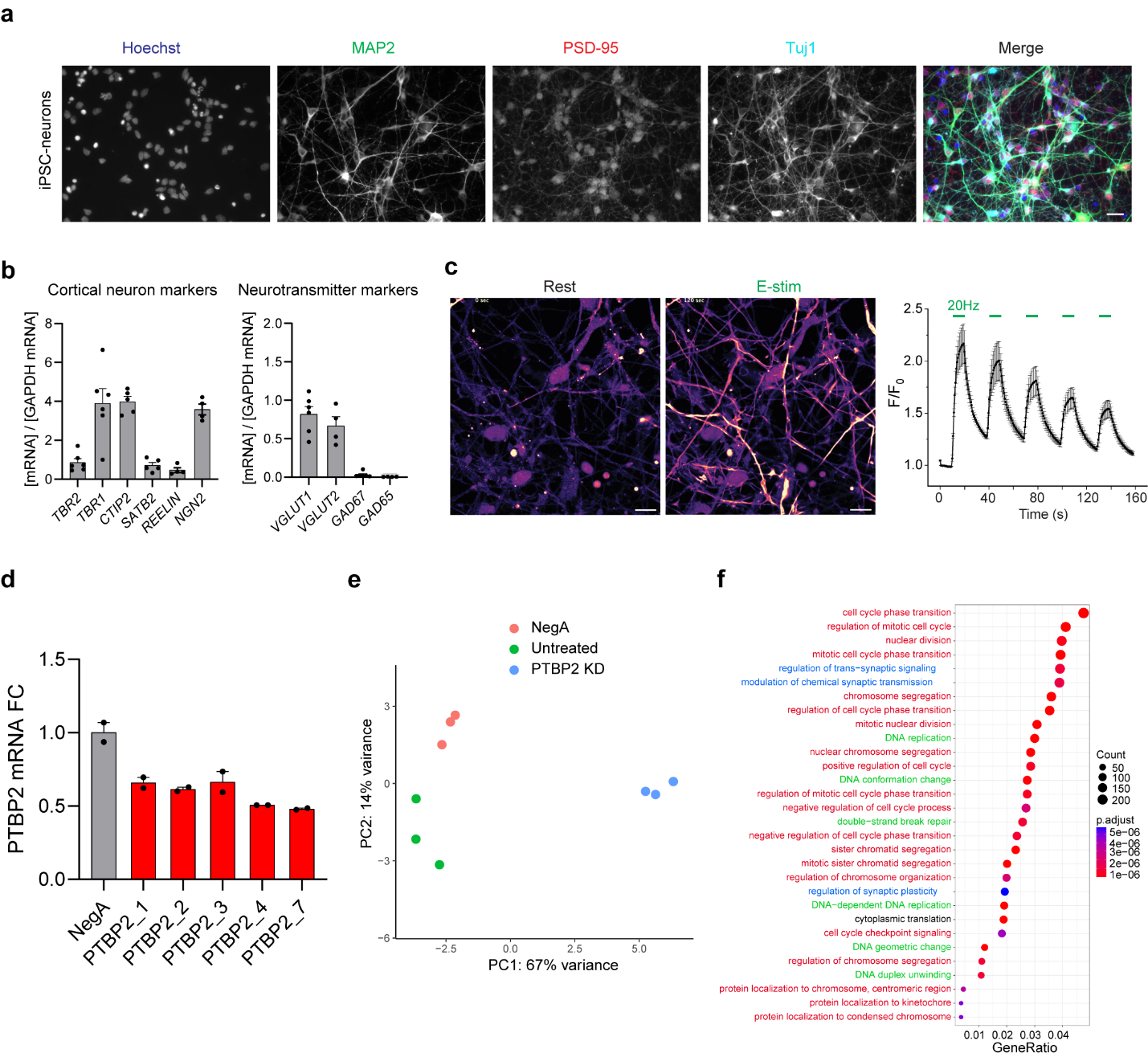

**Supplementary Figure 1. Validation of iPSC-derived cortical neurons and analysis of PTBP2 knockdown in HEK293T cells and iPSC-neurons.** (**a**) Neuronal marker expression by immunofluoresence of iPSC-neurons. iPSC-neurons were immunostained with antibodies specific to MAP2 (green), PSD-95 (red), and Tuj-1 (cyan), and counterstained with Hoechst-33342 (blue), indicative of neuronal phenotype. Scale bar = 20 µm. (**b**) (Left) qPCR of iPSC-neurons for transcripts of the excitatory cortical progenitor TBR2, and cortical neuron markers TBR1, CTIP2, SATB2, REELIN, and NGN2 showing expression of all subtypes, with the highest level transcripts being TBR1, CTIP2, and NGN2. (Right) qPCR of iPSC-neurons identified transcripts of excitatory markers, VGLUT1 and VGLUT2, and minimal expression of inhibitory markers, GAD67 and GAD65. (**c**) Left panel, fluorescence images of iPSC-neurons loaded with a calcium indicator dye (Fluo-4 AM) at rest and electrically stimulated (E-stim) with 20 Hz trains of depolarizing field stimuli lasting 10 s, with 20 s of rest between trains. Scale bar = 10 µm. The right-hand panel shows the quantification of fluorescence intensity changes over time when normalized to initial fluorescence levels (F/F0) using the 20 Hz stimulation. **(d)** qPCR from HEK293T cells transfected with 25 nM of PTBP2 gapmers for 24 h. A non-targeting gapmer (NegA) was included as negative control. **(e)** Principal component analysis (PCA) of gene-level rlog-transformed normalized count data from RNA-seq iPSC-neurons samples. **(f)** Dotplot (clusterProfiler) showing the top results from Gene Ontology (GO) enrichment analysis of genes differentially expressed upon PTBP2 KD (Biological Process, PTBP2 KD vs. untreated iPSC-neurons) relative to a background of all genes evaluated. Gene ratio is number of differentially expressed genes (padj < 0.05) relative to total genes in GO group (Count). In **b**-**d,** data are represented as mean values ± SEM. All data points represent independent biological replicates. **b** (*n* = 4-6 from 3-4 independent differentiations of the CHOP WT10 line). **c** (*n* = 4). **d** (*n* = 2). **e-f** (*n* = 3 biological replicates).

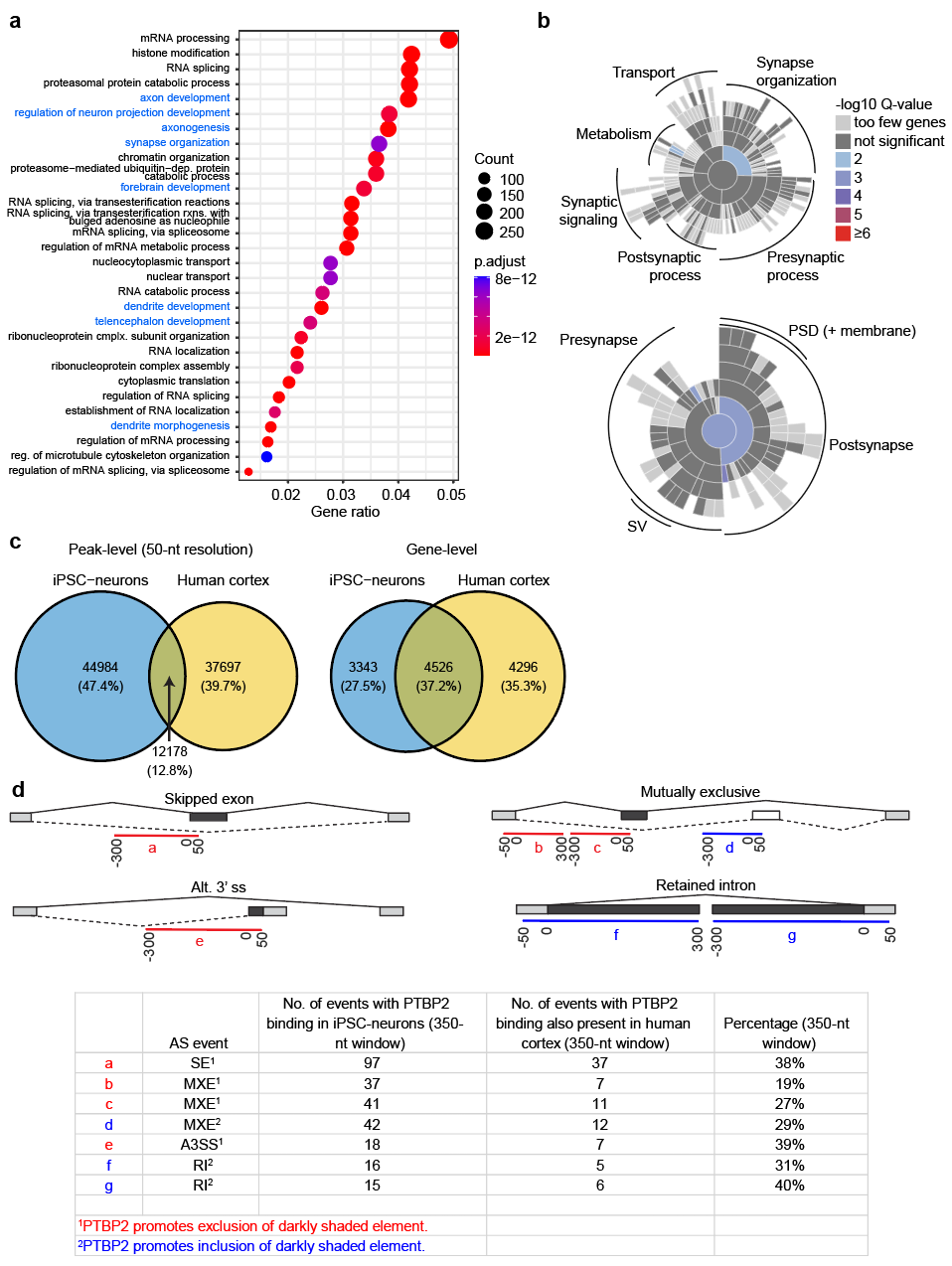

**Supplementary Figure 2. PTBP2 CLIP-seq target gene ontology**. **(a)** Dotplot (clusterProfiler) showing the top 30 categories from Gene Ontology (GO) enrichment analysis (Biological Process) of PTBP2 CLIP-seq peaks in iPSC-neurons. Gene ratio is number of genes with peak calls relative to total genes in Gene Ontology group (Count). Synapse-related terms are highlighted in blue. **(b)** SynGO enrichment analysis of PTBP2 CLIP-seq peaks in iPSC-neurons relative to a background set of brain-expressed genes represented as a sunburst plot. (Top) Biological Process, 442 genes. (Bottom) Cellular Component, 572 genes. **(c)** Venn diagram of PTBP2 binding in iPSC-neurons and human cortex (BA9) at the (left) 50-nt resolution level of CLIP-seq peak calls and (right) at the gene level. **(d)** PTBP2 binding within 350 nucleotides proximal to AS events in iPSC-neurons. (Top) schematic of AS events showing positioning of 350-nt windows (see **Fig. 3f**). (Bottom) Percentage of events with PTBP2 binding in the given 350-nt window in iPSC-neurons that also showed PTBP2 binding in the same window in human cortex (Brodmann Area 9). *n* = 3 biological replicates for PTBP2 CLIP-seq and size-matched input controls.

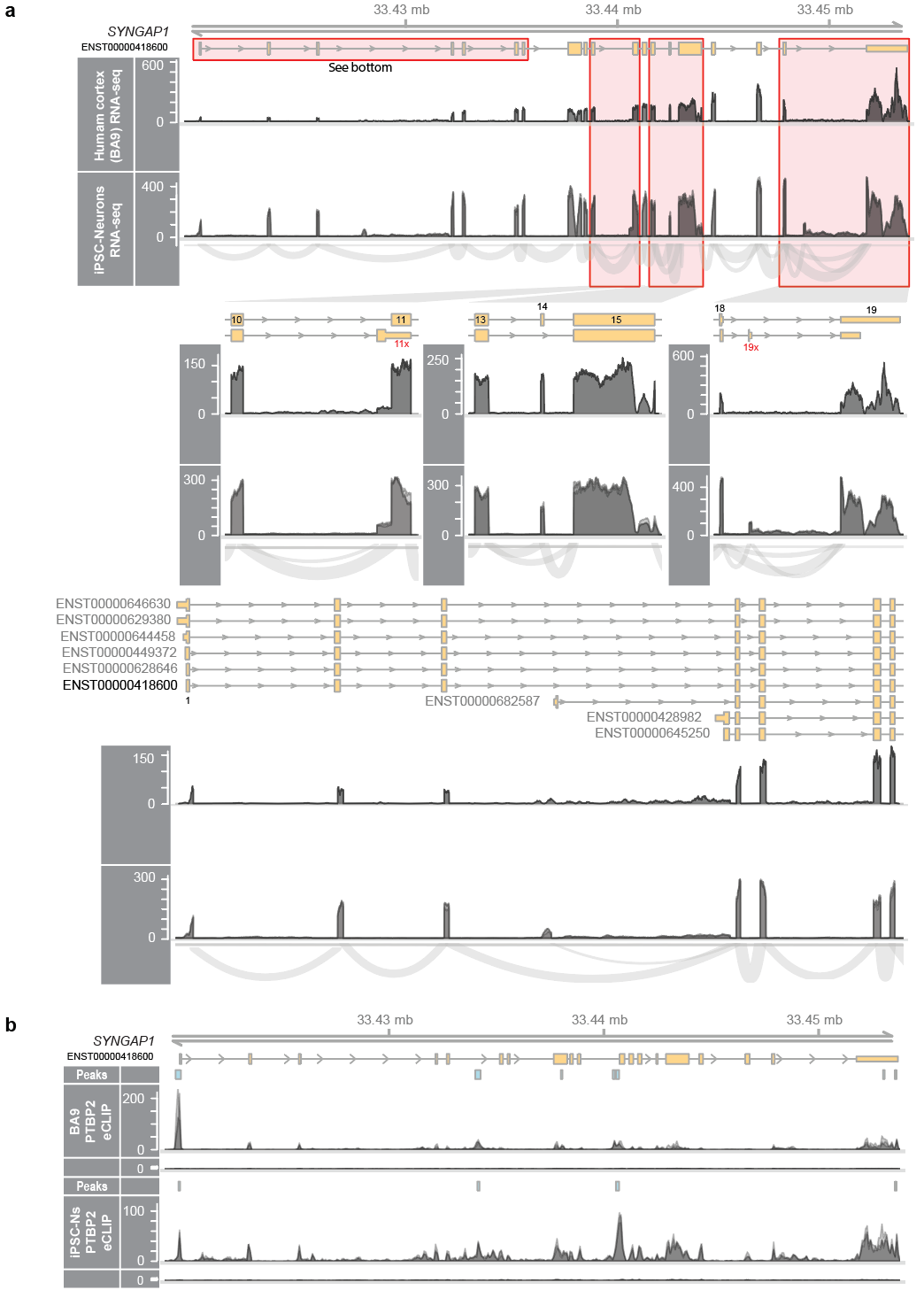

**Supplementary Figure 3. PTBP2 binding and alternative splicing of SYNGAP1**. **(a)** (Top) Gene model of SYNGAP1 followed by RNA-seq read coverage for human cortex (BA9) and RNA-seq read coverage and sashimi plots for untreated control iPSC-neurons. (Insets) Zoom-ins for regions of interest. The human cortex RNA-seq represents 101 samples from GTEx Brain Front Cortex (BA9). *n* = 3 replicates (overlaid) for iPSC-neurons. **(b)** Gene model of ENST00000418600, the dominant SYNGAP1 isoform in brain, followed by human cortex (BA9) CLIP-seq (peaks, PTBP2 eCLIP read coverage, size-matched input read coverage), then iPSC-neurons CLIP-seq (as for cortex). *n* = 3 replicates (overlaid) for PTBP2 CLIP-seq and size-matched input controls.

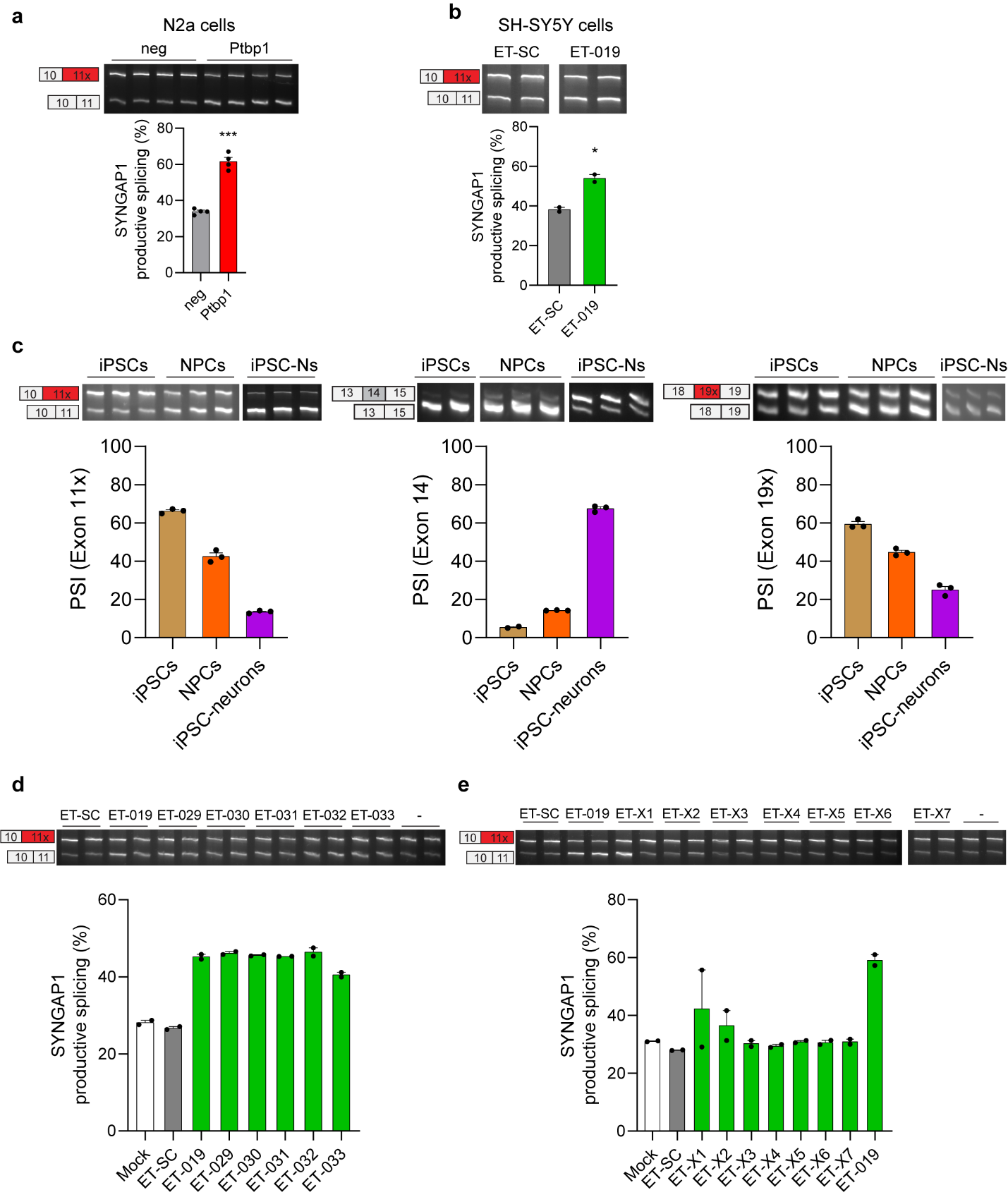

**Supplementary Figure 4. Disrupting PTBP binding in SYNGAP1 site 1 improves SYNGAP1 productive splicing**. (**a**) RT-PCR from N2a cells transfected with either *Ptbp1* gapmer or negative control (neg). (**b**) RT-PCR from SH-SY5Y cells electroporated with 20 µM of ET-019 or negative control ASO (ET-SC) for 24 h. (**c**) Quantification (RT-PCR) of changes in SYNGAP1 AS on (left) exon 11x, (middle) exon 14 and (right) exon 19x in iPSCs, NPCs and iPSC-neurons. (**d**) and (**e**) RT-PCR from HEK293T cells transfected for 24h with 100 nM of ASO targeting SYNGAP1 site 1, including a positive ASO control (ET-019), a non-targeting ASO control (ET-SC) and no ASO control (Mock, -). Data are represented as mean values ± SEM. All data points represent independent biological replicates. **a** (*n* = 4). **b**, **d**, **e** (*n* = 2). **c** (*n* = 3). In **a** and **b,** Student’s t-test. *p < 0.05 and ***p< 0.001.

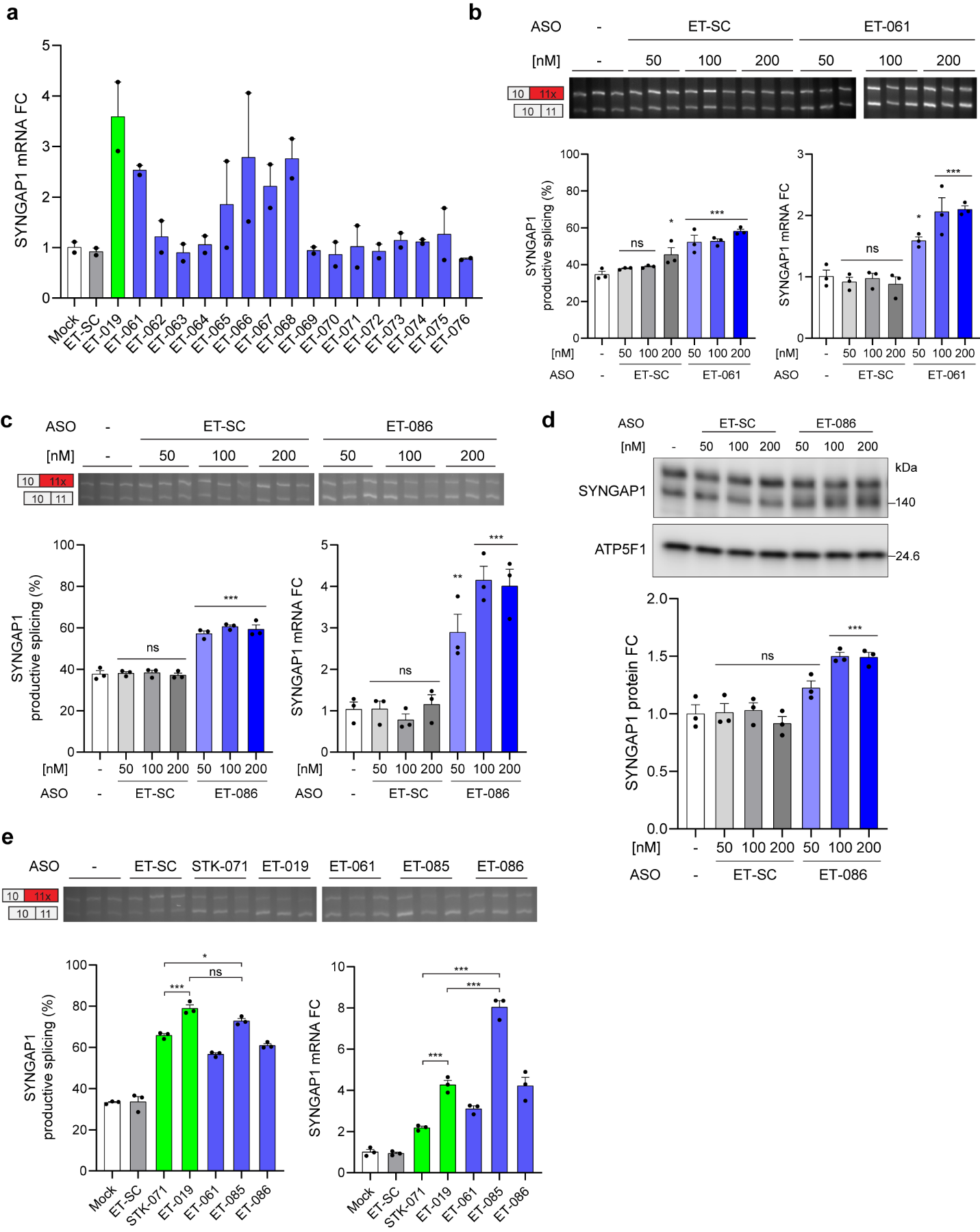

**Supplementary Figure 5. Disrupting PTBP binding in *SYNGAP1* exon 11x upregulates SYNGAP1.** (**a**) qPCR showing SYNGAP1 mRNA levels from samples in Fig. 6B. (**b**) and (**c**) Top and left panels: RT-PCR from HEK293T cells transfected with increasing concentrations of ET-061 (**b**) or ET-086 (**c**) for 48 h, including matching concentrations of the non-targeting ASO control (ET-SC) and no ASO control (Mock, -). Right panels: qPCR showing SYNGAP1 mRNA levels. (**d**) Western blot from HEK293T cells transfected as in (c). (**e**) Top and left panel: RT-PCR from HEK293T cells transfected with 100 nM of ASO for 48 h, including matching concentrations of the non-targeting ASO control (ET-SC) and no ASO control (Mock). Right panel: qPCR showing SYNGAP1 mRNA levels. Data are represented as mean values ± SEM. All data points represent independent biological replicates. **a** (*n* = 2). **b**-**e** (*n* = 3). In **b**-**d**, one-way ANOVA with Dunnett’s multiple comparison test vs mock-treated cells (-). In **e**, one-way ANOVA with Tukey’s multiple comparison test. ns p > 0.05, *p < 0.05, **p < 0.01 and ***p< 0.001.

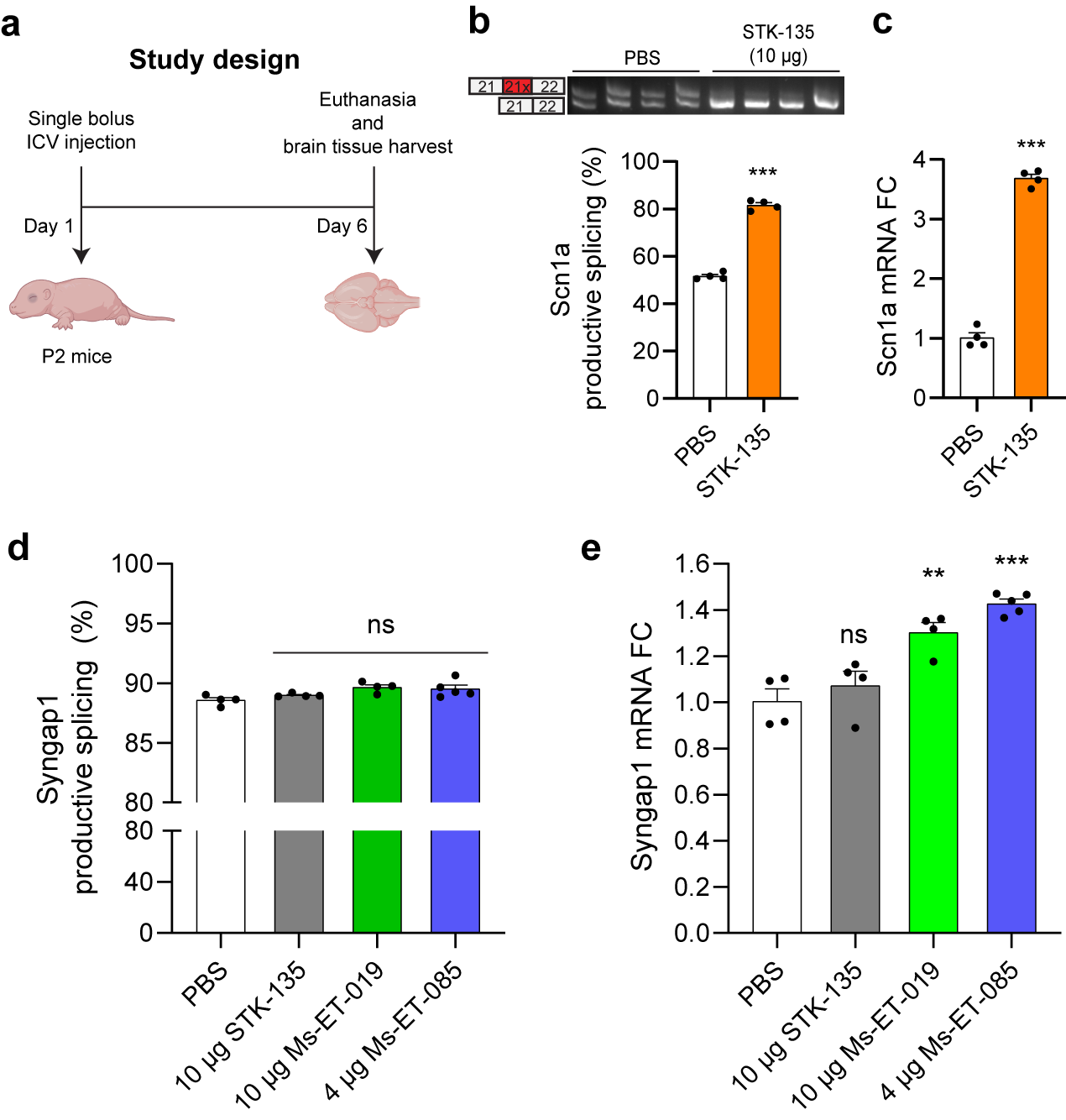

**Supplementary Figure 6. Intracerebroventricular injection of ASOs in neonatal mice increases Syngap1 mRNA expression**. (**a**) Experimental design for evaluation of *Syngap1* ASOs *in vivo*. P2 mice were injected with PBS, a positive control ASO targeting the non-productive exon inclusion in *Scn1a* (STK-135, 10 µg) previously reported by Lim and coworkers^1^, Ms-ET-019 (10 µg) and Ms-ET-085 (4 µg). Mice were euthanized at P7, and brain tissues were harvested and analyzed for productive exon exclusion in *Scn1a* upon STK-135 treatment, and productive exclusion of the alternative 3’ss in *Syngap1* upon *Syngap1* ASO treatments. (**b**) *Scn1a* RT-PCR assay from mouse brains injected with 10 µg of STK-135. The percentage of exon 21x exclusion (productive splicing) in *Scn1a* transcript was calculated based on densitometric analysis of RT-PCR products. (**c**) qPCR showing *Scn1a* productive transcript levels. (**d**) RT-PCR assay of *Syngap1* Ex11 productive splicing. (**e**) qPCR showing *Syngap1* productive transcript levels. Data are represented as mean values ± SEM. All data points represent independent biological replicates. **b**, **c** (*n* = 4). **d**, **e** (*n* = 4 except *n* = 5 for Ms-ET-085). In **b** and **c**, Student’s t-test. In **d** and **e**, one-way ANOVA with Dunnett’s multiple comparison test vs PBS. ns p > 0.05, **p < 0.01 and ***p< 0.001.
