## Supplementary material for "Mapping PTBP splicing in human brain identifies targets for therapeutic splice switching including *SYNGAP1*": Description of Supplementary Files

**Description of Additional Supplementary Files**

File Name: Supplementary Movie 1

Description: iPSC-neurons loaded with a calcium indicator dye (Fluo-4 AM) at rest and electrically stimulated with 20 Hz trains of depolarizing field stimuli lasting 10 s, with 20 s of rest between trains.

File Name: Supplementary Data 1

Description: Differential gene expression analysis in iPSC-neurons following PTBP2 KD.

File Name: Supplementary Data 2

Description: Table of differentially spliced genes in iPSC-neurons (PTBP2 KD vs. Untreated) along with Orphanet status, up-regulation/down-regulation at the gene level, and CLIP-seq peak calls proximal to splice event.

File Name: Supplementary Data 3

Description: Sequence and chemistry of oligonucleotides used in study.

File Name: Supplementary Data 4

Description: Primer sequences used for assessing expression of neuronal markers.
